# Protein turnover in the developing *Triticum aestivum* grain

**DOI:** 10.1101/2021.06.15.448508

**Authors:** Hui Cao, Owen Duncan, A. Harvey Millar

## Abstract

Protein abundance in cereal grains is determined by the relative rates of protein synthesis and protein degradation during grain development. Through combining *in vivo* stable isotope labelling and in-depth quantitative proteomics, we have measured the turnover of 1400 different types of proteins during wheat grain development. We demonstrate that there is a spatiotemporal pattern to protein turnover rates which explain part of the variation in protein abundances that is not attributable to differences in wheat gene expression. We show that approximately 20% of total grain ATP production is used for grain proteome biogenesis and maintenance, and nearly half of this budget is invested exclusively in storage protein synthesis. We calculate that 25% of newly synthesized storage proteins are turned over during grain development rather than stored. This approach to measure protein turnover rates at proteome scale reveals how different functional categories of grain proteins accumulate, calculates the costs of protein turnover during wheat grain development and identifies the most and the least stable proteins in the developing wheat grain.

## Introduction

Wheat (*Triticum aestivum* L.) is a trusted source of protein and calories for human consumption and serves as the staple food for 30% of the human population^1^. Although global wheat production continues to grow steadily, it is not sufficient to meet predicted demand, especially when the global population is projected to exceed 9 billion by 2050 requiring an increase of wheat production by about 70%^2, 3^. To cope with such a challenge, researchers, breeders and wheat-processing industries have been working collaboratively to enhance yield while still maintaining grain-quality attributes such as grain protein content^4, 5^.

While wheat protein content is dominated by glutenin and gliadin storage proteins^6^, it also includes thousands of other components spread through the endosperm, embryo and pericarp^7, 8^. The abundance of high and low molecular weight glutenins determine dough elasticity, while gliadins contribute to dough extensibility^9^. A complex pattern of expression of the protein synthesis apparatus and many different types of proteases occur during the early and later stages of grain development^10-12^. Further, wheat storage proteins are assembled, folded, aggregated and stabilised in the ER, and then follow a Golgi or ER route to vacuolar protein bodies^13, 14^. As a consequence, the abundance of specific proteins products correlates poorly with expression of the encoding genes in both wheat^15^ and many plants^16, 17^. Identification and quantification of additional factors that contribute to individual protein abundances and changes in abundance profiles over time are needed to better control wheat grain protein composition.

One such factor is protein turnover rate that dictates how quickly new plant proteins are synthesized while unwanted or dysfunctional proteins are degraded and recycled^18-20^. Study of protein turnover rate in plants have focused on either model plants such as Arabidopsis or non-food tissues of crops, such as seedlings or leaves^19, 21-26^. These non-food plant proteomes can be studied in steady-state conditions influenced only by dilution through tissue growth^18^ and the relatively short lag period for introduction of stable isotopes makes calculating protein turnover measurements in these systems feasible^19, 26^. In comparison, studying grain filling is complicated by the complex dynamics of protein accumulation during development and the relative difficulty of rapidly introducing stable isotopes into the spike^27^.

Here, we have overcome these technical and biological challenges to establish *in vivo* stable isotope (^15^N) labelling to enable protein turnover rates to be measured during wheat grain development. Using this approach, we monitored synthesis and degradation rate of over 1400 different wheat protein types during grain filling and calculated the ATP energy usage for protein synthesis and degradation. Wheat grain protein turnover rates provide scientists and breeders with a quantitative and in-depth understanding of how each wheat grain protein accumulates and a biotechnological pathway to craft a lower cost grain proteome.

## Results

### In *vivo* stable-isotope nitrogen (^15^N) labelling of wheat grain proteins

Measuring the synthesis and degradation of individual wheat proteins requires the labelling of new proteins in grains. We undertook this labelling by incorporation of the stable isotope of nitrogen, ^15^N, into the amino acids used for their synthesis^18, 20^. Wheat grain development includes cell division and expansion until 14 days post anthesis (DPA), grain filling (14 to 28 DPA), and desiccation and maturation (28 to 42 DPA)^28^. To determine when during development the turnover rate of wheat grain proteins could be measured, four sets of wheat plants were raised in hydroponic medium with N salts of natural isotope abundance (99.6% ^14^N, 0.4% ^15^N) until plants reached 7, 14, 21 and 28DPA, respectively. The growth media of natural N isotopes was then replaced by heavy ^15^N labelled media (2% ^14^N, 98% ^15^N). Grain samples were harvested after 7 days of continuous labelling (Extended Data Fig. 1a). Peptide mass spectrometry analysis showed that nearly 25% of N atoms in newly synthesised proteins in wheat grain were ^15^N after the 7-day labelling period, regardless of the grain DPA (Extended Data Fig. 1b). Broadly, calculations of the labelled protein fraction can be made independently of the % of ^15^N they contain, making this technique robust to differences in ^15^N incorporation rate^18-20^. We found younger grains showed a higher proportion of newly synthesised proteins in their total protein pool (labelled protein fraction, LPF) than older grains (Extended Data Fig. 1c). However, we know that at least 20% of ^15^N incorporation is required to make high-quality protein turnover rates measurements with FDR < 1%^26^, we worked to achieve this basal rate by focusing on 7DPA-old grain as our first time point for subsequent labelling experiments. To capture the major events of grain development, samples of progressively labelled grain were collected after 3, 7 and 10 days of continuous ^15^N labelling to span the 7DPA to 17DPA period (Extended Data Fig. 2a). A significant expansion was seen in grain size over this time period with single grain fresh weight tripling over 10 days from 25 mg at 7DPA to 78 mg at 17DPA (Fig. 1a, b). The ^15^N enrichment level gradually increased over time from 20% after 3 days (10DPA) to 29% after 10 days (17DPA) (Fig. 1c).

**Fig. 1.**
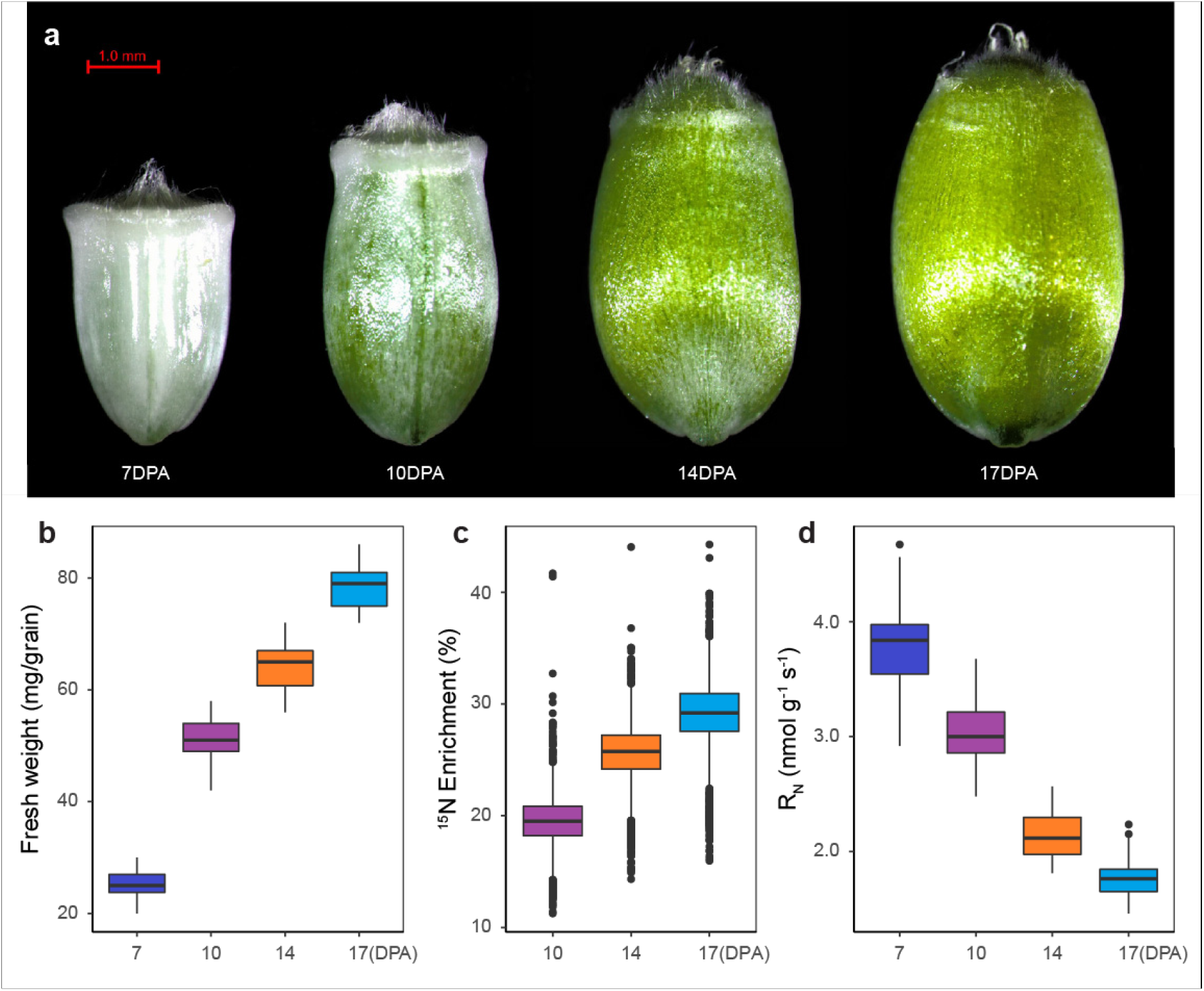
Time-dependent changes in grain size, fresh weight, respiration rate and ^15^N labelling of newly synthesised protein. **a**, A representative image of wheat grain at different developmental stages. **b**, The fresh weight of individual grain at different developmental stages (*n* = 64). **c**, The ^15^N enrichment in newly synthesised proteins at 10, 14 and 17DPA after a switch to ^15^N media at 7DPA. **d**, Respiration rate (R_N_) of wheat grain at different developmental stages expressed as O_2_ consumption rate (nmol O_2_ per gram fresh weight per second; *n* = 64). Detailed data used in this analysis are listed in Supplementary Data 1.

### One day lag time in labelling of wheat grain proteins

Incorporation of nitrogen into the developing grain requires long-distance transport from the roots through the phloem, before its assimilation, use and storage in leaves and the spike^29^. Using a logarithmic regression model and our 3, 7 and 10 day labelling data, we calculated a lag of 28 hours (Extended Data Fig. 3a). To confirm this lag time estimated, we used both GC-MS analysis of free amino acids and LC-MS analysis of peptides to assess the ratio of heavy (+1) to mono abundance of amino acids and peptides in samples collected in the first 30 hours of labelling. We found both amino acids and peptides of proteins remained at their natural abundance level before 24 h but rapidly rose between 24-30 h following ^15^N labelling (Extended Data Fig. 3b, c).

### Grain respiration rate and ATP production

Protein production is a major user of cytosolic ATP generated through oxidative phosphorylation. Using a fluorophore-based oxygen sensor, we measured the grain respiration rate during grain development and found younger grains respired at twice the rate of older ones on a fresh weight basis (Fig. 1d). As cellular respiration is the primary ATP production source in wheat grain, and assuming a cellular ATP production ratio of 1 O_2_ to 4.5 ATP^19^, the total ATP production rate of a single grain was calculated to be 37, 60, 54 and 54 µmol ATP per day for grains at 7, 10, 14 and 17DPA, respectively (Supplementary Data 1d).

### Calculating individual fold changes in protein abundance for proteins during wheat grain development

As the proteome of the wheat grain is not in a steady-state during filling, we needed to obtain individual fold changes in protein abundance (FCP) values for each protein of the wheat grain. To do this a spike-in procedure was developed that involved adding a standard amount of a fully ^15^N labelled wheat grain standard (see methods) to four grains from each time point. This helps to define proteins rapidly accumulating in the grain and exclude the effect of differential starch accumulation on the measurements (Extended Data Fig. 2b). Using this approach, quantification of the abundance change of 2307 non-redundant grain proteins were measured using data from over 71,000 independently quantified peptides (Supplementary Data 4a). Principal component analysis of the quantitative data set showed that grain samples separated according to their DPA with the first principal component explaining nearly 94% of the variation. Only minor variation was observed in the second principal component and biological replicate samples of the same time point were closely clustered together (Extended Data Fig. 5a). Over half of the identified proteins increased in abundance ≥ 2-fold from 7DPA to 17DPA, with a few exceptions that showed the opposite pattern, consistent with net protein accumulation during grain development (Fig. 2a). Quantitative abundance changes for different proteins during grain development varied from 0.018-fold to 126-fold. Examples shown in Fig 2b illustrate the dynamics of different protein groups such as ribosomal proteins (fold change < 3 over 10 days) and storage proteins (fold change > 25 on average and up to 126 over 10 days).

**Fig. 2.**
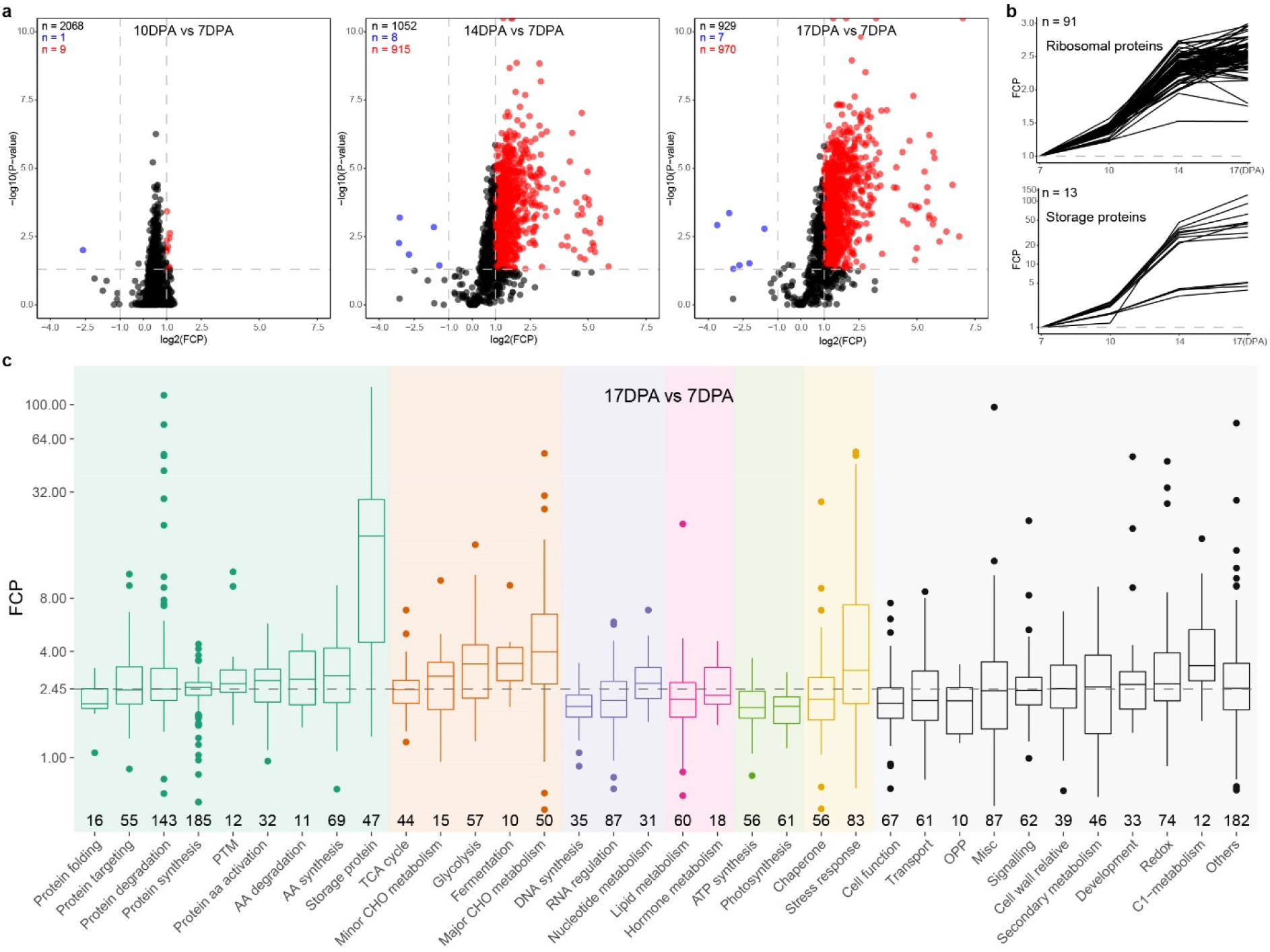
Fold change in abundance (FCP) of 2307 proteins during wheat grain development. **a**, Volcano plots of all fold changes in protein abundance at each time point. Proteins with a fold change in abundance ≥ 2 or ≤ 0.5 and with a p-value ≤ 0.05 are shown by red circles (increase abundance) or blue circles (decrease abundance), respectively. All other data points are show as black circles. The number of proteins of each colour is shown on the top-left corner of each plot, and dashed lines indicate the cut-off value of fold change in protein abundance and p-value. **b**, Examples of slow FCP of ribosomal proteins compared to the rapid accumulation of storage proteins during grain development. **c**, Proteins with fold change in abundance between 7DPA and 17DPA are shown in 34 functional categories (≥ 10 proteins) within 7 broad functional categories. Proteins in all other categories (≤ 10 proteins per category) are grouped into the ‘Others’ category. The number of proteins in each category is displayed along with x-axis, and functional categories were sorted within each super category by increasing median FCP. The y-axis is log2 transformed FCP and the dashed line shows the overall median FCP across all wheat grain proteins. Pairwise t-test results between categories are listed on Supplementary Data 4b. Broad functional categories are: Amino acid metabolism (Green), Carbohydrate metabolism (Orange-red), Nucleotide metabolism (Purple), Lipid metabolism (Pink), Energy producing (Lawngreen), Stress response (Gold) and ‘Other’ categories (Black).

MapMan functional category bins^30^ were used to separate identified proteins into 34 groups (≥ 10 proteins in each). These showed a median fold change value of 1.38-, 2.31- and 2.45 at 10, 14 and 17DPA compared to 7DPA (Fig. 2c and Extended Data Fig. 6). Exceptions were found in the categories of storage proteins, major CHO metabolism and stress response, where fold change values significantly higher than the median were observed. These data are consistent with biogenesis and accumulation of starch and storage proteins being major events during grain development and moisture loss in ripening and maturation being a post-anthesis stress^31^. Protein categories associated with protein folding, TCA cycle, DNA synthesis and photosynthesis showed lower fold change than the median, likely because they were less active cellular pathways during grain development. Together this evidence showed clear temporal patterns in the regulation of protein abundance during grain development.

### Calculating wheat grain protein synthesis and degradation rates during grain development

The labelled protein fraction (LPF) is an instantaneous measurement of the ratio between existing and new protein molecules for each individual proteins of interest at a given point of time. This information is derived from the ratio of peptides for a protein with greater than natural abundance (labelled) to the sum of peptides with natural abundance and greater than natural abundance [^15^N/(^14^N + ^15^N)]. These measurements are derived from quantitation of the deviation of the isotopic envelope from what would be expected of natural abundance peptides of a particular sequence^19, 32^. LC-MS/MS results from 252 fractionated samples, including 108 fractions of progressive labelling samples and 144 fractions of unlabelled samples, provided data on over 44,000 quantified peptides. Combining the information from peptides derived from the same protein allowed us to measure the LPF of 1711 non-redundant proteins from wheat grains. The median of the relative standard deviation from the mean (RSD) for each protein at a given time point was less than 11% across all three time points (Supplementary Data 5a). Spearman correlation coefficient analysis showed a relatively high correlation of LPF between biological replicates ranging from 0.81 to 0.95 (Extended Data Fig. 5g). PCA analysis suggested that up to 97% of the variation observed could be explained by the first principal component alone (Extended Data Fig. 5d). Through combining the individual fold change in protein (FCP) and LPF values for specific proteins, we then calculated protein synthesis rates and degradation rates for 1447 non-redundant proteins (Supplementary Data 5b). Relative protein synthesis rates (K_S_/A; where K_s_ is the synthesis rate constant, and A the abundance of protein) varied nearly 100 fold; ranging from 0.057 d^-1^ to 5.27 d^-1^ with a median of 0.35 d^-1^. By contrast, protein degradation rates (K_D_; where K_D_ is the degradation rate constant) had a median of 0.11 d^-1^ and ranged from effectively zero up to 0.94 d^-1^. The latter indicates a range of proteins in grain with half-lives of less than one day.

To determine if subcellular location influenced protein turnover rates in wheat grain, we grouped the data set by their subcellular location retrieved from a database of crop proteins with annotated locations; cropPAL^33^. We reassigned storage proteins into vacuole because once assembled in ER, storage proteins are transported to storage protein vacuoles (PSV)^34^. Based on this, vacuole proteins, predominantly storage proteins, showed higher median synthesis and degradation rates than other subcellular structures. The same pattern was found in subcellular protein sets of the peroxisome, Golgi apparatus and extracellular secreted proteins (Et), but the opposite pattern was seen in protein sets located in the mitochondrion, cytosol, plastid and the endoplasmic reticulum (ER) (Fig. 3a). These observations generally agreed with published data from Arabidopsis leaves which showed organellar proteins that are physically separated from the cytosolic proteolysis system have relatively lower degradation rate, while cellular structures such as the Golgi apparatus have a frequently updated protein complement and a relatively higher turnover rate^19^.

**Fig. 3.**
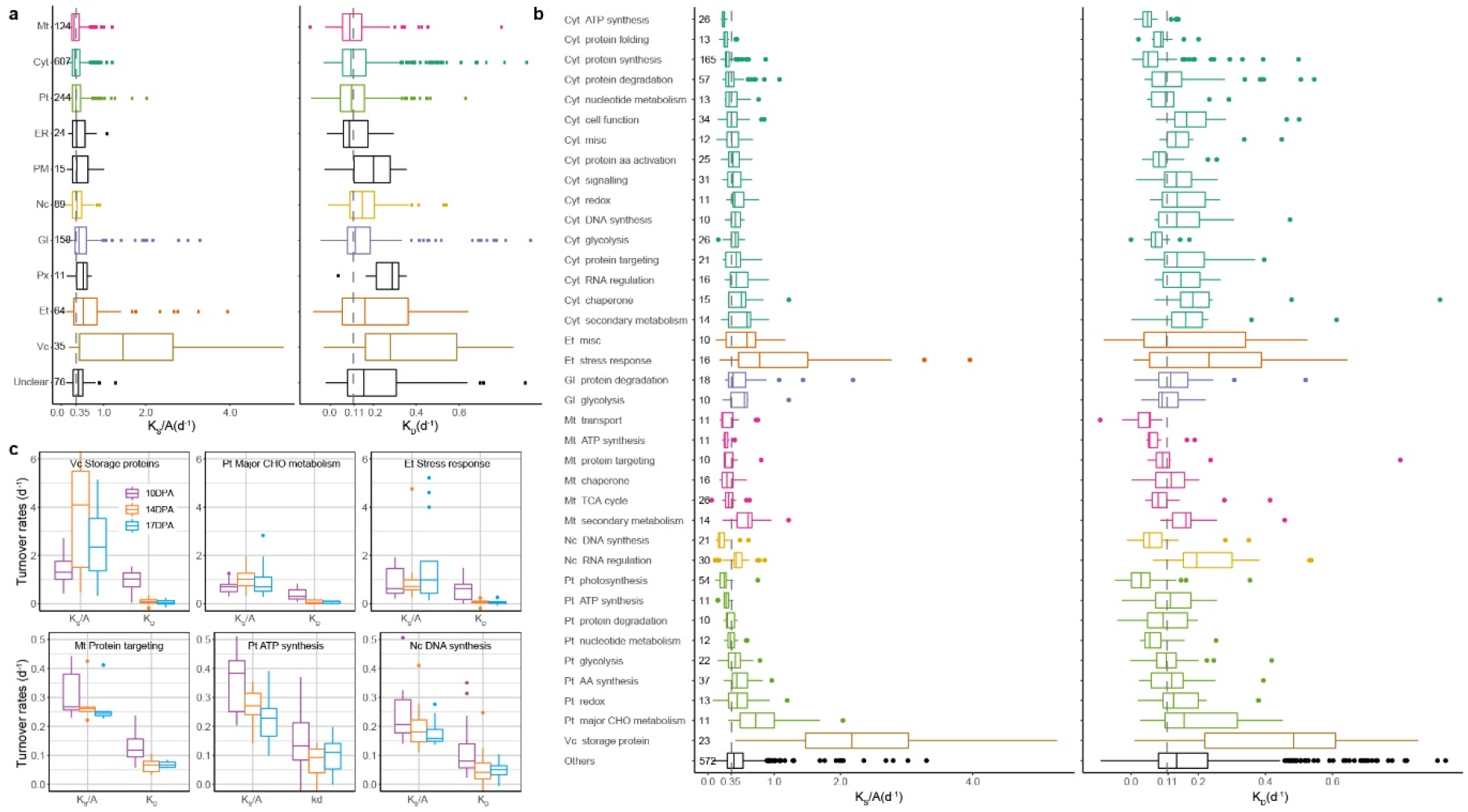
Rates of synthesis and degradation of 1447 wheat grain proteins during grain development. **a**, The averaged protein synthesis (K_S_/A) and degradation (K_D_) rates calculated across all three time points grouped by the subcellular location of each protein (CropPAL.org). The major subcellular locations highlighted are Mitochondria (Mt, Pink), Cytosol (Cyt, Green), Plastid (Pt, Lawngreen), Nucleus (Nc, Gold), Golgi Apparatus (Gl, Purple), Extracellular (Et, Orange-red), and Vacuole (Vc, Peru). Dashed lines indicate the overall median K_S_/A and K_D_ across all proteins. The number of proteins in each subcellular location is displayed next to the y-axis. Boxes are sorted by increasing order of median K_S_/A. **b**, The averaged protein synthesis and degradation rates calculated across the three time points grouped by both subcellular location and functional category. Only major functional categories with protein no. ≥ 10 were displayed, and minor categories with protein no. < 10 were grouped into ‘Others’. The colour scheme was the same used in **a**. **c**, Examples of 6 functional categories of interest showing the largest difference (upper panel) and smallest difference (lower panel) between K_S_/A and K_D_ at each time point. Detailed statistical analysis is explained, and relevant results are listed in Supplementary Data 5c.

When the data were arranged by MapMan functional category bins, most of the 37 major groups maintained a median K_S_/A of 0.35 ± 0.15 d^-1^ and median K_D_ of 0.11 ± 0.05 d^-1^ (Fig. 3b). However, exceptions were found, notably stable photosynthesis proteins had a median K_S_/A of 0.24 d^-1^ and median K_D_ of only 0.03 d^-1^, while storage proteins had a median K_S_/A of 2.17 d^-1^ and median K_D_ of 0.48 d^-1^, indicating rapid synthesis and degradation cycling of a portion of storage proteins in developing grains. Variation of protein turnover rates was also observed for proteins located in the same organelle or subcellular structure but which belonged to different functional categories, or vice versa, such as cytosolic proteins involved in ATP synthesis, protein folding and protein synthesis, or RNA regulation factors located in the cytosol and nucleus (Fig. 3b). To better understand the connection between protein functional roles and their turnover rates, we compared turnover rate profiles between three sets of rapidly-cycling proteins and three more stable house-keeping protein sets. The K_S_/A of the rapidly-cycling proteins gradually increased or stayed steady over development, while a much lower synthesis rate and decreasing K_S_/A over development was observed for more stable house-keeping functional categories (Fig. 3c). As expected, much lower K_D_ was determined for house-keeping proteins in comparison with rapidly-cycling proteins, but unlike protein synthesis rate, both protein types shared a similar pattern over development, notably that K_D_ dropped from its peak at 10DPA to its lowest level at 14DPA followed by a slight increase by 17DPA.

Storage proteins were prominent members of lists of the 20 most rapidly synthesized proteins (55%) and the 20 most rapidly degraded proteins (30%) in wheat grains; this included gliadins, globulins, high-molecular-weight glutenin subunits (HMW-GS) and low-molecular-weight glutenin subunits (LMW-GS) (Extended Data Table 1). Proteins annotated as responding to stress conditions were the second most numerous in both lists; 4 were on the 20 most rapidly synthesized list and 3 were on the most rapidly degraded list. An alpha-amylase inhibitor (TraesCS6D01G000200.1) topped the degradation rate list and had a calculated half-life of only 0.74 d^-1^.

To facilitate further comparisons, we used filters to select a set of 149 wheat proteins that were synthesized rapidly (K_S_/A ≥ 2*median K_S_/A of 0.71 d^-1^) and a set of 67 proteins that were synthesized slowly (K_S_/A ≤ 0.5*median K_S_/A of 0.18 d^-1^). These two groups represented roughly 10% and 5% of all measured proteins, respectively (Supplementary Data 5b). Adopting the same filtering principles, 291 (∼20%) and 320 (∼22%) proteins were selected as rapidly and slowly degrading proteins, respectively. These groups were then further clustered as being house-keeping proteins (ie. both slow K_S_/A and K_D_, SS), induced but stable proteins (i.e. fast K_S_/A and slow K_D_, FS), and rapidly-cycling proteins (ie. both fast K_S_/A and K_D_, FF) (Extended Data Fig. 7a). Functional category analysis of these sets showed that the house-keeping proteins mainly participated in photosynthesis, DNA synthesis and glycolysis; while induced but stable proteins participated in major CHO metabolism, transport and amino acid synthesis proteins; and the rapidly-cycling protein set was dominated by storage proteins, proteins involved in stress response and proteins involved in protein degradation (Extended Data Fig. 7b-d).

### Protein turnover rates explain varying abundance of proteins in different parts of the grain

To find connections between protein turnover rates and the spatiotemporal abundance patterns of grain proteins, we separately quantified the abundance of 5550 grain protein groups in endosperm, embryo and pericarp (Extended Data Fig. 8 and Supplementary Data 6). These proteins were then assigned into seven categories based on their protein abundance profile using the definition and equations explained in the Methods section. Forty percent of these proteins were found evenly balanced in their abundance across grain tissues, ∼32% were assigned as tissue-specific proteins (7% for endosperm, and 12.5% for embryo and pericarp, respectively) and ∼29% as tissue-suppressed proteins, i.e. proteins present two tissues but low in the third (14% for endosperm, and 7.5% for embryo and pericarp, respectively) (Fig. 4a). The endosperm-specific proteins (Fig. 4c) had high fold changes and increasing net accumulation rate over development, while pericarp-specific proteins (Fig. 4c) had low fold change in abundance and decreasing in net accumulation rate over time. In comparison, embryo-specific proteins had an increasing net accumulation rate over development but very low turnover rates and fold change in abundance values (Fig. 4c). The proteins with tissue-balanced profiles, showed moderate fold changes and net accumulation rates remained stable over development (Fig. 4c). This pattern was also observed to a lesser extent in protein groups suppressed in a specific tissue (Extended Data Fig. 9).

**Fig. 4.**
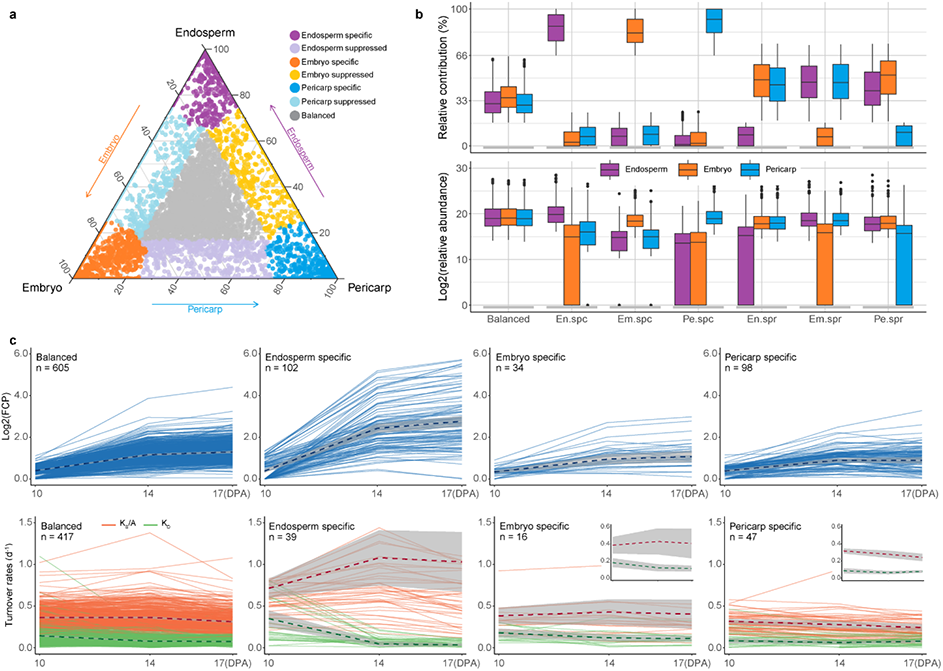
Changes in protein abundance and protein synthesis and degradation rate profiles of proteins expressed in different grain tissue types during grain development. **a**, A ternary plot of the abundances of 5550 protein groups measured in Endosperm, Embryo and Pericarp extracts from grain. Each circle represents a protein and its position indicates the relative contribution of each protein to grain tissue in grain protein abundance. Proteins with a relative contribution ≥ 66% in one tissue and a protein concentration ≥ 3-fold that in the other two tissues are defined as tissue-specific proteins (shown close to triangle vertices), proteins with a relative contribution ≤ 17% in one tissue are defined as tissue-suppressed proteins (between vertices and close to edges), while the remaining proteins are found relatively evenly balanced across grain tissues (grey circles in the middle). **b**, Box plots of the relative protein contribution (upper panel) and actual relative abundance estimated via label free quantification (lower panel) for each tissue in each protein expression category. **c**, Line plots showing FCP (Blue), protein synthesis rate (Orange) and protein degradation rate (Green) of balanced and tissue specific protein sets during grain development. Only proteins having values (FCP or turnover rates) at all three time points are included. The dashed line demonstrates the mean values, and the grey shade area shows the 95% confidence intervals. The optimized y-axis scale version are inserted for embryo and pericarp specific proteins to highlight the change pattern.

### ATP Energy budget for wheat grain proteome synthesis and maintenance

The ATP energy cost for both synthesis and degradation of 1140 proteins in the wheat grain proteome was calculated through combining Intensity Based Absolute Quantification (iBAQ) values (Supplementary Data 7a), protein turnover rates, amino acid length, and the ATP cost per amino acid residue for protein synthesis and protein degradation^35-37^ (Supplementary Data 7b). This revealed that during grain development, each grain invested 20% of its total ATP production on protein biogenesis and maintenance. The ATP budget for protein synthesis (9.8 µmol grain^-1^ d^-1^) was nearly 9 times larger than for protein degradation (1.2 µmol grain^-1^ d^-1^; Fig. 5a).

**Fig. 5.**
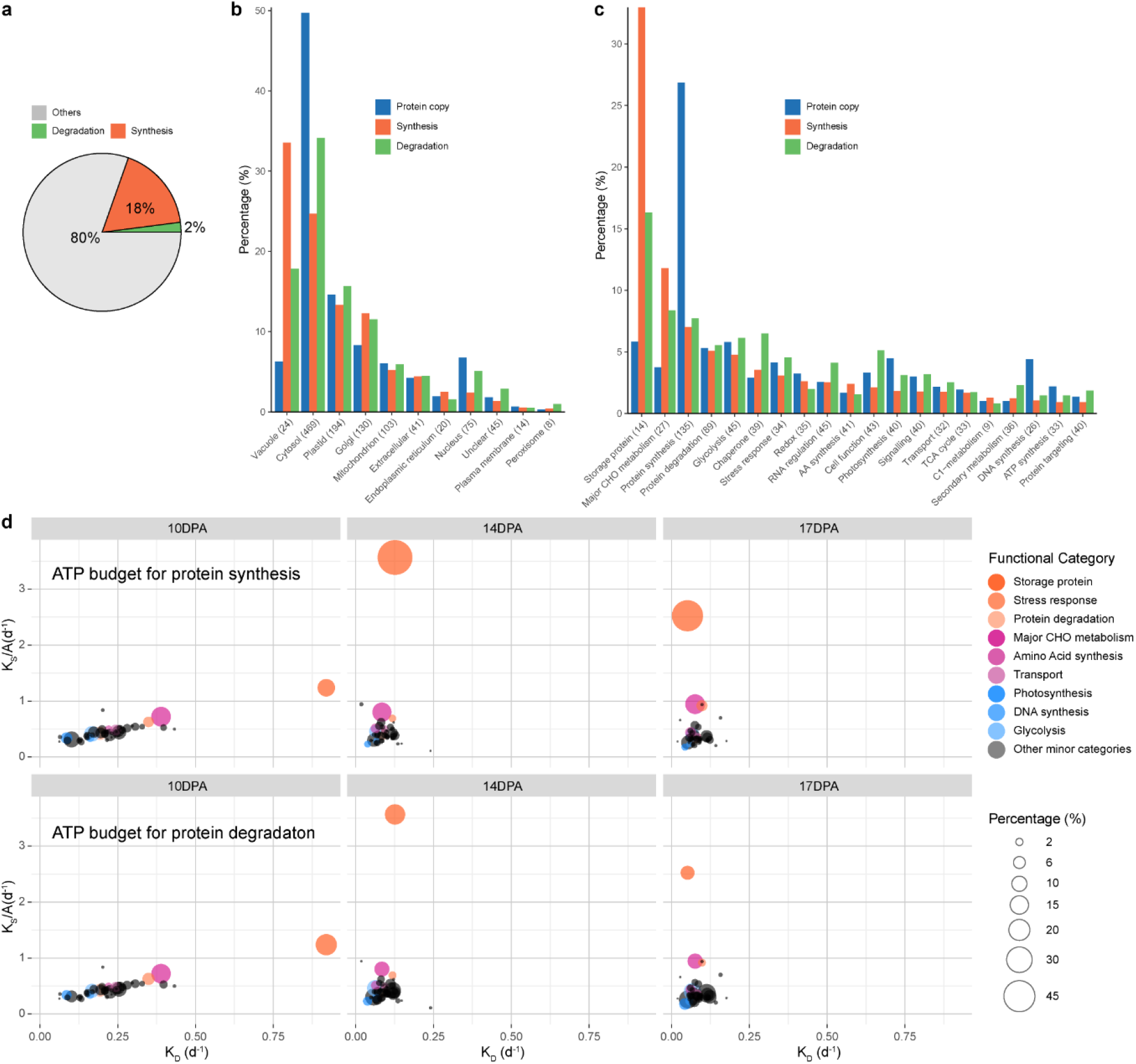
ATP energy budget used in wheat grain proteome synthesis and maintenance during grain development. **a**, The overall proportion of cellular ATP budget used for protein synthesis, protein degradation, and other cell events and maintenance. **b**, proportional distributions of protein copy numbers (iBAQ) and ATP cost of protein building and maintenance of major cellular organelles and subcellular structures. Proteins in location groups sorted by decreasing ATP usage for protein synthesis. The number of proteins in each category is included within brackets. **c**, proportional distributions of protein copy numbers (iBAQ) and ATP cost of protein building and maintenance of protein in major functional categories. **d**, Bubble plots showing, by the size of circles, the changing profiles of cellular ATP energy budget for different classes of proteins during grain development. The top three functional categories having both fast K_S_/A and K_D_ rates (Orange-red), fast K_S_/A and slow K_D_ rates (Violet-red), and both slow K_S_/A and K_D_ rates (Blue) are highlighted. Calculations are based on grain total ATP production in **a**, but based on total ATP energy budget for protein synthesis or degradation in **b, c** and **d**.

Grouping the ATP usage data by the subcellular location of each protein allowed us to assess the relative energy cost of maintaining different subcellular structures in the wheat grain. Maintaining the proteins of the vacuole (the destined location of storage proteins) used one third of the total ATP budget for protein synthesis although this group only contained 24 unique groups of proteins. This was followed by the largest category, cytosolic proteins (comprising 469 protein groups), that cost ∼25% of the ATP, and the plastid and Golgi apparatus each cost ∼13% of the ATP used for protein synthesis (Fig. 5b). The cytosol proteome cost 34% of the total ATP for protein degradation, which was nearly twice that of the vacuole (17.8%). Expressing the ATP usage rate as nmol per million protein copies per grain per day (nmol ATP mcopy^-1^ grain^-1^ d^-1^), the vacuole proteome was the most expensive for protein synthesis on a location basis, costing 144 nmol ATP mcopy^-1^ grain^-1^ d^-1^, which was 15 times higher than the lowest cost compartment, the nucleus, at only 9.3 nmol ATP mcopy^-1^ grain^-1^ d^-1^. Peroxisomes topped the list of protein degradation energy costs, consuming 12 nmol ATP mcopy^-1^ grain^-1^ d^-1^, while the cytosol, by contrast, only consumed 2.2 nmol ATP mcopy^-1^ grain^-1^ d^-1^ for this purpose.

When the data was grouped by functional category, we observed that storage proteins and enzymes of major CHO metabolism were the most costly in terms of both protein synthesis and degradation, together they used half the energy budget of total protein synthesis and a quarter of the total energy budget for protein degradation (Fig. 5c). Storage proteins were the most expensive functional category in terms of synthesis and maintenance of proteins, costing 152 nmol ATP mcopy^-1^ grain^-1^ d^-1^ for synthesis and additional 9 nmol ATP mcopy^-1^ grain^-1^ d^-1^ for degradation. Less costly protein synthesis and maintenance expense was evident for ribosomal proteins (7 nmol ATP mcopy^-1^ grain^-1^ d^-1^ for synthesis and 0.9 nmol ATP mcopy^-1^ grain^-1^ d^-1^ for degradation). As expected, the 20 highest cost proteins were predominately storage proteins and major CHO metabolism proteins, among which the highest energy cost for an individual protein was an 11S globulin (TraesCS1A01G066100.1) that cost nearly 2% of grain ATP in its production during this stage of grain development (Extended Data Table 2).

Collectively, storage proteins dominated the wheat grain proteome in terms of abundance, fold change in protein, protein turnover rates and ATP energy usage; but this prominence builds over time. The grain invested only 10% of its total ATP energy budget for protein synthesis on the formation of storage proteins at the early grain filling stage (10DPA), but this subsequently grew 4.5 times in the next 4 days, reaching 45% of the total ATP budget for protein synthesis by 14 DPA, and settled at 36% by 17DPA (Fig. 5d). The ATP energy budget for degradation of storage proteins showed a similar pattern, increasing at the early stage of grain filling and decreasing at later stages, although it remained at a relatively high level over time. The machinery of major CHO metabolism, which is needed for starch synthesis, showed a rather stable energy usage over time compared to storage proteins. Likewise, proteins in functional categories such as photosynthesis, DNA synthesis and glycolysis showed a relatively stable energy usage over time and only consumed a minor proportion of ATP production in the grain (Fig. 5d).

### Wheat storage proteins accumulation during grain filling

To illustrate the complexity of accumulation patterns of members of the seven key storage protein families and their causes from 7DPA to 17DPA we integrated 6 different measures into a single radar graphic (Fig. 6a). Our findings indicated that except for globulin 1 encoding genes which peaked at 30DPA, the mRNA of key storage proteins were most abundant at the early grain filling stage and were temporally offset from protein accumulation. A common pattern in synthesis rates and biogenesis energy cost (CostSyn) was shared between storage protein families; they reached the highest level at 14DPA and remained at this level until at least 17DPA. In comparison, significantly higher degradation rates and energy consumption were measured at the pre-grain filling stage (10DPA) than the grain filling stage (from 14DPA). Collectively these features resulted in a rapid accumulation in storage protein abundance, which increased over 20-fold on average and up to 90-fold for some storage protein isoforms from 10DPA to 17DPA (Fig. 6b).

**Fig. 6.**
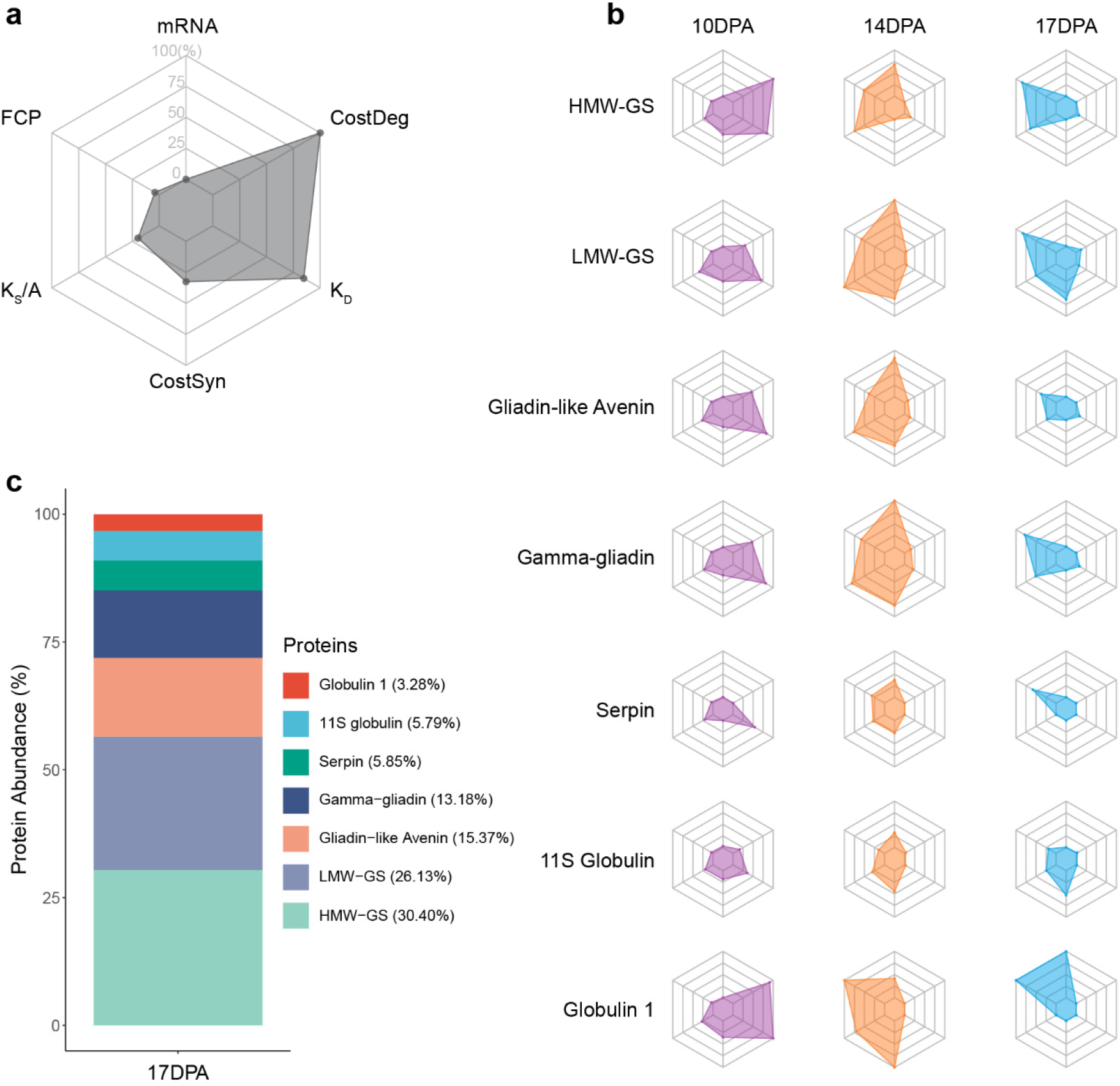
Accumulation profiles of key storage proteins during grain filling. **a**, The example radar chart. Six categories of data were collected and analysed, namely transcript data (mRNA, rang: 0-1000 tpm), fold change in protein abundance (FCP, rang: 1-25 folds), protein synthesis rate (K_S_/A, rang: 0-5.84 d^-1^), ATP energy cost for protein synthesis (CostSyn, rang: 0-290.2 nmol ATP mcopy^-1^ grain^-1^ d^-1^), protein degradation rate (K_D_, rang: 0-1.36 d^-1^), and ATP energy cost for protein degradation (CostDeg, rang: 0-51.6 nmol ATP mcopy^-1^ grain^-1^ d^-1^). mRNA ≥ 1000 tpm and FCP ≥ 25-fold were treated as 100%, while missing values were replaced by 0 for visualization. The percentage normalisation was conducted across all proteins and time points allows comparisons to be made in time series and between protein types. The raw data and statistical test results are collected at Supplementary Data 8. **b**, Radar charts showing the six molecular profiles of the key wheat grain storage proteins at 10, 14, and 17DPA during grain filling. **c**, Abundance of the key wheat grain storage proteins at 17DPA.

## Discussion

In this study, we provide an extensive and quantitative understanding of how different wheat grain proteins accumulate during grain development. The resulting protein turnover data allows the costs of individual proteins, protein functional groups, and cell structures during grain development to be calculated, providing an innovative foundation for grain protein engineering strategies.

Previous studies calculated a lag time of 8 h for ^15^N labelling of proteins in hydroponically growth barley leaves^26^ and 5 h in Arabidopsis young leaves^19^, but the lag involved in long-distance transport from roots to the wheat spike is likely to be considerably longer^29^. Furthermore, most protein turnover studies in plants to date have assessed steady-state scenarios where there is little or no developmental change in protein composition^19, 26, 38^, which is not the case in grain filling. We solved these two problems by experimentally calculating a 28 h lag time for ^15^N incorporation (Extended Data Fig. 3) and measuring the fold change in abundance for each protein of interest during grain development using a spike-in of a fully ^15^N labelled proteome (Fig. 2 and Extended Data Fig. 6). This has allowed not only the protein turnover rates of over a thousand proteins to be determined, but to do so in a background of proteins changing in relative abundance over a 0.018-fold to 126-fold range.

As autotrophs, plants have access to a large source of ATP via photosynthesis in source leaves. However, the ATP yielded by this process is largely invested in sucrose synthesis that is transported to sink tissues, which leaves oxidative phosphorylation by respiration as the primary source for cytosolic ATP-dependent processes like protein synthesis and degradation^19, 39^. Based on this we calculated the proportional energy cost for synthesis and degradation of the wheat grain proteome during grain filling based on grain respiration rates (Fig. 5). Our data show that approximately 20% of total grain ATP production through respiration is used for grain proteome biogenesis and maintenance, and nearly half of this budget is invested exclusively in storage protein synthesis. This proportion was similar in size to reports of the cost of Arabidopsis leaf protein turnover which varied from 16-42% of total ATP depending on the age of the examined leaf^19^. However, in grains this ATP investment is co-commitment with the high ATP demand of starch synthesis at the same times during development^40, 41^. Collectively, ATP provision to protein synthesis and degradation and the starch synthesis represent an energy constraint that will dictate the relative investment in starch and protein in the final grain.

Dough rheological properties and bread-making quality in wheat are primarily determined by gluten quantity^9^. In our development analysis, four storage protein families that are key components of gluten polymers, namely HMW-GS, LMW-GS, gliadin-like avenin and gamma-gliadin, represent 85% of storage protein abundance (Fig 6c and Supplementary Data 8); consistent with previous studies reporting 80-90% of total grain protein is gluten polymers^6, 42^. Extensive previous studies have also demonstrated that gluten quality, both the ratio of gliadin/glutenin and the ratio of HMW-GS/LMW-GS, strongly influences dough viscosity and elasticity, thus governing bread-making quality traits^43-45^. The ratio of HMW-GS/LMW-GS is typically ∼0.67 in mature grains^42^, our data show HMW-GS/LMW-GS continuously decreased overtime from 3.66 at 10DPA to 1.63 by 14DPA and ending up 1.16 by 17DPA. Unlike the ratio of HMW-GS/LMW-GS, the ratio of gliadin/glutenin remained stable during grain development in our hands (Supplementary Data 8). Protein turnover data reveals that 25% of these newly synthesized storage proteins appear to be turning over during grain development rather than being stored and that instability of storage proteins was most prominent early in grain development between 10 and 17 DPA. Wheat storage proteins are generally assembled, folded and aggregated in the endoplasmic reticulum (ER), followed by a trafficking event mediated by protein bodies to vacuoles for storage through either a Golgi-vacuole route or the ER-vacuole route^13, 14^. Previous evidence has suggested that storage proteins are stabilized during the processing within the ER due to the formation of complex polymers through disulphide and noncovalent bonds (inter-and/or intra-chain bonds)^14, 46^. Dominguez and Cejudo showed that extractable wheat grain endoprotease activity reached its maximum at 15 DPA and then decreased afterwards^10^. Likewise, Nadaud et al showed grain proteolysis activity peaked at approximately 10DPA, and showed a decrease pattern over the following 4 days^11^. Our data also indicate a significant decrease in protein degradation machinery of the endomembrane system from 10DPA to 17DPA (Supplementary Data 7c). These high degradation rates of storage proteins detected at 10DPA are not the case for all storage protein isoforms (Fig. 6b and Supplementary Data 7c), providing a breeding or biotechnological pathway to select for lower cost storage proteins with lower degradation rates.

Extensive effort has been placed into assessing the correlation between transcript level and protein abundance, however only ∼40% of the variation in protein abundance is typically explained by transcript data in eukaryotes, including plants^16, 17, 47, 48^. In wheat there is only a 32% concordance between protein and transcript expression profiles observed during grain development^15^. This atlas of wheat grain protein turnover rates contributes to explaining the remaining variation in protein abundance not correlated to transcript levels. As an atlas iy shows that despite variation of turnover rate for different proteins approaching 100 fold, protein turnover rates are highly correlated with their changing profiles of FCP over time when a net accumulation rate calculation is considered as the key driver of protein abundance alterations (Fig. 3 and Extended Data Fig. 10). It is also known that spatiotemporal regulatory mechanisms broadly exist in live-systems and play essential roles in protein abundance regulation to meet various biological scenarios^19, 49-51^. This pattern is also evident in our wheat grain data and we show that the higher turnover rates of storage proteins when compared to other protein groups drives the preferential accumulation of storage proteins as one of the largest cellular events during grain filling (Fig. 2-4).

Overall this study provides a comprehensive and in-depth view of which key storage proteins accumulation during pre- and the early-mid grain filling stage. In addition, the quantitative data for non-storage proteins provides insights into a wide range of biological events during wheat grain development, such as the protein machinery for starch accumulation and the initiation of desiccation stress responses. Future studies will be needed to reveal why wheat grains partake in cycles of protein synthesis and degradation and find approaches to alter this cycling such as knockout of unstable storage proteins and expression of more stable versions using either conventional cross breeding approaches or gene editing technologies. Success in this pursuit could help increase grain protein content, improve energy use efficiency of protein production, and help meet the unprecedented food demand for sustainable plant-based protein production in modern agriculture.

## Methods

### Plant materials and growth condition

Wheat (*Triticum aestivum*) plants of cv. Wyalkatchem, were grown in an indoor chamber under 16/8-h light/dark conditions with 26/18°C, 60% humidity and light intensity of 800 µmol m^-2^ s^-1^. The hydroponic growth system used for plant growth is described by Munns and James^52^. Fifty litres Hoagland solution was used for 24 plants and changed weekly^8^. Seeds were vernalized under 4°C for 3 days before being transferred to the growth chamber followed by 3 to 5 days germination until two leaves emerged at which point they were transferred to the hydroponic growth system. Plants were firstly grown with natural abundance N medium, which was then replaced with ^15^N (^15^NH_4_ NO_3_, 98 %, Sigma fine chemicals) medium for a labelling programme when plants reached the required growth stage (eg. 7DPA). Plants were rinsed three times with distilled deionized water before being transferred to the ^15^N medium. A set of fully labelled plants to generate the spike-in standard samples were grown with ^15^N medium at all times after germination. Grain samples, including unlabelled, progressive labelled and fully labelled samples, were harvested at multiple time points as shown in Extended Data Fig. 1 and 2. Only 8 grains from the middle of the ear (both floret 1 and 2) were collected in each case. Grain tissues of embryo, endosperm and pericarp were hand-dissected by scalpel. Due to the challenge of separating embryo from endosperm in young grain, only 14 and 17DPA-old grain were used for dissection and tissue samples of two time points were pooled together. Samples were snap frozen in liquid nitrogen and stored in −80°C for further protein extraction.

### Grain respiratory oxygen consumption rate measurement

Respiration rates of single premature grain at different growth age (7, 10, 14, and 17DPA) were measured using a Q2 oxygen sensor (Astec-Global) in sealed 2 mL capacity tubes at 24°C. The O_2_ concentration within tubes were measured at an interval of 5 min for 16 h. In total, 64 replicates, including 8 biological replicates (from different plant) with 8 technical replicates (8 grains from the same ear), were performed for each growth stage. The O_2_ consumption rate (R_N_) trace was generated using a moving slope of O_2_ consumption in a 2 h window and the formula reported by Scafaro et al in 2017^53^. The representative R_N_ was calculated using the O_2_ consumption slope in the 2 h window from 1-3 h. Total ATP production of a single wheat grain at each growth stage was also estimated based on the ATP production rates of 1 O_2_ to 4.5 ATP^19^.

### Spike-in approach for individual wheat grain FCP measurement

To precisely measure individual FCP of wheat grain proteins during grain development, a spike-in approach was developed. Briefly, an equal number of ^15^N fully labelled grains (8 grains in this study) of each time points (7, 10, 14 and 17DPA) were pooled together and ground using mortar and pestle under liquid nitrogen. The fully labelled fine sample powder of 100 mg was spiked into four unlabelled grains at each time point as an internal standard, followed by another grinding (Extended Data Fig. 2b). These spike-in containing samples were store in −80°C for subsequent protein extraction, mass spectrometry data acquisition and FCP measurement.

### Sample preparation

A chloroform/methanol extraction protocol^54^ was applied for total protein extraction in this study. Briefly, 200 mg sample powder generated by extensive grinding in liquid N_2_ was mixed with 400 μL extraction buffer (125 mM Tris-HCl pH 7.5, 7% (w/v) SDS, 0.5% (w/v) PVP40, Roche protease inhibitor cocktail (Roche, 1 tablet per 50 ml)) and rocked on ice for 10 min. After a centrifugation at 10,000 g for 5 min, about 200 μL supernatant was transferred into a new 2ml eppendorf tube, followed by protein precipitation through mixing the supernatant with 800 μL methanol, 200 μL chloroform and 500 μL distilled deionized water. The pellet was washed twice using methanol and then incubated with 90% (v/v) acetone at −20 °C twice for at least 1 hour each time. After drying at room temperature, protein pellet was resuspended using resuspension buffer (50 mM Ambic, 1% (w/v) SDS and 10 mM DTT). Protein concentration was quantified by an amido black method^55^.

Proteins (200 μg) were incubated with 20 mM DL-dithiothreitol for 20 min in darkness at room temperature, followed by a second incubation with 25 mM iodoacetamide for 30 min in darkness at room temperature. After diluting the SDS to its working concentration at 0.1% (w/v) via adding distilled deionized water, proteins were digested overnight using trypsin (Promega, Sequencing Grade Modified Trypsin, USA) at 37°C with protein-trypsin ratio of 50:1. The SDS removal and high-pH, reversed phase peptide fractionation for digested peptide solution^56^ were conducted on an off-line HPLC (1200 series, Agilent Technologies) combining with two J4SDS-2 guard columns (PolyLC) and an XBridge^TM^ C18 3.5 μm, 436 x 250 mm column (Waters). The pump flow was set at 0.5 ml/min using the following solution B (90% acetonitrile with 10 mM ammonium formate (pH 10/NH_4_OH) gradient: 2 to 5% in 6 min, 5 to 35% in 60 min, 35 to 70% in 13 min, 70 to 100% in 10 min and 100 to 2% in 3 min. In total, 64 fractions for each sample were collected from 15 to 79 min in 1 min windows. The first 12 and last 4 fractions were discarded and the rest of the fractions in the same column of the 96 well plate were combined together. The final 12 fractions of each sample were dried down through a vacuum centrifuge and store in −80°C for further mass spectrometry analysis.

### LC-MS data acquisition and processing

Peptide fractions were resuspended with 25 μL of 5% (v/v) acetonitrile and 0.1% (v/v) formic acid in HPLC grade water followed by a filtering step using 0.22 μm centrifugal filters (Millipore). Purified peptide suspensions (2 μL each) were injected into a HPLC-chip (Polaris-HR-Chip-3C18, Agilent Technologies) through a capillary pump with a flow at 1.5 μL/min. Peptides were eluted from the C18 column online into an Agilent 6550 Q-TOF. Gradients were generated by a 1200 series nano pump (Agilent Technologies) with the nano flow at 300 nl/min, of which 5-35% (v/v) solution B (0.1% (v/v) formic acid in acetonitrile) in 35 min, 35-95% in 2min and 95-5% in 1 min. Parameters setting in MS acquisition was as described previously^8^. In total, LC-MS data of 516 fractions were successfully collected (Extended Data Fig. 2d). The primary MS data files are available via ProteomeXchange with identifier PXD022231.

To measure individual FCP of wheat grain proteins during grain development, LC-MS data stored in Agilent .d files of 288 fractions of spike-in and unlabelled samples (144 fractions each) were first converted to mzML files using the online Trans Proteomic Pipeline (TPP, v.5.2.0)^57^. The Comet search of above mzML files was conducted against protein database from IWGSC (v.1.0, 137029 sequence)^58^ using decoy search, 20 ppm peptide mass tolerance and maximum 2 missed cleavage^59^. Further PeptideProphet search was performed for each replicate at each time point (eg. 12 fractions of replicate 1 of spike-in data at 7DPA and 12 fractions of replicate 1 of unlabelled data at 7DPA) and results were merged into single analysis file using PPM to accurate mass binning and decoy hits to pin down negative distributions. Protein identification and corresponding LPF for each protein were obtained by using an in-house pipeline written in R, of which 12 mzML files of spike-in data for each replicate were mapped back to its corresponding PeptideProphet file (.pep.xml file). Filters of probability > 0.8, FDR < 3%, rsd ≤ 25% or sd ≤ 25% of overall mean LPF were applied for peptide list, while filters of probability > 0.95, FDR < 1%, rsd ≤ 25% or sd ≤ 25% of overall mean LPF, independent identifiers ≥ 3 across all samples, and total quantified peptides ≥ 4 were used in protein list. As LPF is the ratio of heavy nitrogen to total nitrogen H/(H+L), the relative abundance for each protein in the ratio of heavy to light (H/L) was estimated by (1-LPF)/LPF. Individual FCP of each protein was calculated using the relative abundance at each time point dividing by its value at 7DPA. The same strategy as the FCP dataset with further LPF ≥ 0.2 was applied to progressive labelling dataset to obtain LPF values of each protein.

The LC-MS data of unfractionated 30-h labelling samples (Extended Data Fig. 2) were processed through Agilent MassHunter Workstation (v.10.1) to determine the ratio of heavy (+1) to mono abundance of peptides. Peptide peak area was used as peptide relative abundance.

### GC-MS data acquisition and processing

Grain sample powder of 30 mg were mixed with 250 μL metabolite extraction buffer (85% (v/v) methanol, 15% (v/v) distilled deionized water, and 0.1mol L^-1^ sorbitol (D-Sorbitol-^13^C_6_) as the internal standard), and subsequently incubated on a thermomixer at 75°C and 950 rpm for 10 min. After mixing with 125 μL of chloroform and 250 μL of distilled deionized water, sample solutions were centrifuged for 15 min at 2,000g, of which 50 μL of supernatant was transferred to a new tube and dried out using a vacuum centrifuge. Sample derivatization started with an incubation in 20 μL of 20 mg mL^-1^ methoxyamine hydrochloride in pyridine for 2 h at 37°C, followed by a second incubation in 20 μL of *N*-methyl-*N*-(trimethylsilyl)-trifluoroacetamide (SIGMA, USA) for 30 min at 37°C. Incubations were conducted on a thermomixer at 950 rpm. Volume of 40 μL derivatized sample was transferred to glass vials for GC-MS analysis. Metabolites samples of 1 μL were injected into an Agilent 7890A gas chromatograph coupled with a Varian CP9013-Factor 4 column (40 m 3 0.25mm i.d.) and an Agilent 5975 quadrupole mass spectrum detector. GC-MS data acquisition was performed following descriptions reported by O’Leary and co-workers^60^. GC-MS data were processed and analysed using Agilent MassHunter Workstation (v.10.1) as mentioned earlier.

### Label free quantification using MaxQuant

The same unlabelled LC-MS data of 144 fractions were processed by searching against IWGSC database through MaxQuant (v.1.6.1.0, http://www.maxquant.org/) with the iBAQ quantitation algorithm, 20 ppm peptide mass tolerance, maximum 2 missed cleavage and FDR < 1% for absolute protein copy number estimation^61, 62^. Using the same parameter setup and LFQ search algorithm instead of iBAQ, the relative protein abundance of grain tissue samples (including endosperm, embryo and pericarp, 108 fractions in total) were also obtained. The reported protein lists were further filtered with rsd ≤ 25% or sd ≤ 25% of overall mean abundance, independent identifiers ≥ 3 across all samples, and total quantified peptides ≥ 4. The protein contributions of each single of the three tissues to the total protein abundance were calculated using formulas as follows

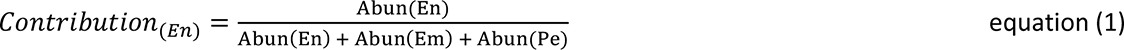

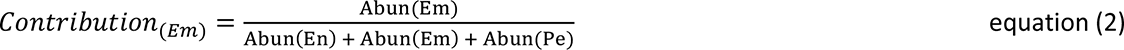

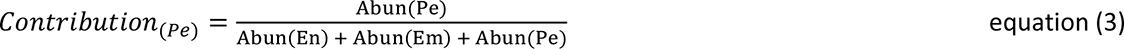

Where En, Em and Pe represent endosperm, embryo and pericarp, and Abun represents the protein relative abundance. Proteins with a relative contribution ≥ 66% in one tissue and ≥ 3-fold that in the other two tissues are defined as tissue-specific proteins (shown close to triangle vertices), while non tissue-specific proteins with a relative contribution ≤ 17% of a tissue are defined as tissue-suppressed proteins. The remaining proteins are found relatively evenly balanced across grain tissues. Protein category definition was further confirmed by the independent Tau method^63^ (Extended Data Fig. 8c). Ternary diagram visualization was performed using the ggtern R package^64^.

### Protein turnover rates and ATP energy cost calculation

Protein synthesis rates and degradation rates were calculated using formulas as follows

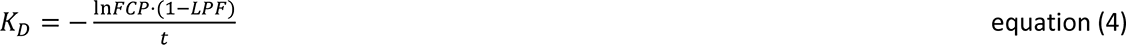

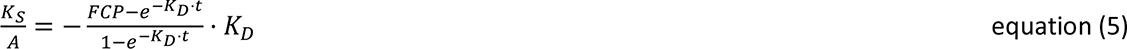

Where A is the protein starting abundance at the beginning of the labelling programme (7DPA). According to previous reports, protein synthesis cost was 5.25 ATP per residue including ribosome translation, protein transport and amino acid biosynthesis^35, 37^, while the cost for protein degradation was approximately 1.25 ATP per residue through proteasome degradation pathway^36^. The ATP energy cost for protein turnover were therefore estimated by combining absolute protein copy number, amino acid length, protein turnover rates, grain ATP production and ATP energy cost per residue. The full-length intact peptide sequences of Chinese Spring glutenin subunits and gliadins were referred to long read sequencing data^65, 66^. Detailed step-by-step explanations of both protein turnover rates and its ATP energy cost calculation were previously reported^19, 67^.

### Gene expression data retrieval

Gene expression data of the key storage proteins were downloaded from the Wheat Expression Browser website (http://www.wheat-expression.com/)68,69. Only Chinese Spring grain tissue data at 2, 14 and 30DPA under non-stress conditions (choulet_URGI) were used in this study.

### Statistical analysis

Data processing, statistical analysis and visualization were performed in the R environment (v.3.5.1). Statistical tests and replicate number are as shown in figure and figure legends.

## Acknowledgements

H.C was supported by Research Training Program Fee Offset – International Student and UWA Safety-Net Top-Up Scholarships. This work was supported by Australian Research Council funding to A.H.M (CE140100008; FL200100057)

## Author contributions

H.C, O.D and A.H.M conceived and designed the project. H.C and O.D performed the experiment and data analysis. H.C wrote the manuscript, A.H.M and O.D read and corrected the manuscript.

## Extended Data Tables

**Extended Data Table 1.** The 20 most rapidly synthesized and degraded wheat grain proteins during grain development.

| Accession | Protein Name | Location | Functional Category | Rates<br>(d<sup>-1</sup>) | Brep<br>No. | Pep<br>No. | Spectra<br>No. | Half-Life<br>(d) |
| --- | --- | --- | --- | --- | --- | --- | --- | --- |
| **20 most rapidly synthesized proteins** |  |  |  |  |  |  |  |  |
| TraesCS1D01G000300.1 | LMW-GS | vacuole | storage protein | 5.27 | 5 | 8 | 15 | NA |
| TraesCS1A01G007400.1 | Gamma-gliadin | vacuole | storage protein | 4.37 | 7 | 7 | 19 | NA |
| TraesCS1D01G000200.1 | LMW-GS | vacuole | storage protein | 4.27 | 6 | 8 | 30 | NA |
| TraesCS1A01G066100.1 | Globulin-11S | vacuole | storage protein | 4.22 | 6 | 27 | 49 | NA |
| TraesCS7B01G072000.1 | Alpah-AI | extracellular | stress response | 3.96 | 7 | 8 | 11 | NA |
| TraesCS1B01G084300.1 | Globulin-11S | vacuole | storage protein | 3.41 | 6 | 13 | 21 | NA |
| TraesCS3A01G095600.1 | Alpha-AI | golgi | stress response | 3.3 | 4 | 4 | 8 | NA |
| TraesCS4B01G328000.1 | Alpah-AI | extracellular | stress response | 3.27 | 4 | 6 | 8 | NA |
| TraesCS1B01G330000.1 | HMW-GS | vacuole | storage protein | 3.16 | 7 | 21 | 32 | NA |
| TraesCS5A01G554200.1 | Beta-amylase | golgi | major CHO metabolism | 3.02 | 6 | 14 | 21 | NA |
| TraesCS7D01G031700.1 | Gliadin-like avenin | vacuole | storage protein | 2.89 | 6 | 14 | 32 | NA |
| TraesCS4A01G296000.1 | Globulin 1 | vacuole | storage protein | 2.85 | 4 | 6 | 8 | NA |
| TraesCS1A01G007700.1 | Gamma-gliadin | vacuole | storage protein | 2.84 | 3 | 5 | 12 | NA |
| TraesCSU01G032700.1 | Beta-amylase | golgi | major CHO metabolism | 2.8 | 8 | 44 | 92 | NA |
| TraesCS4D01G325000.1 | Alpah-AI | extracellular | stress response | 2.77 | 4 | 8 | 22 | NA |
| TraesCSU01G103300.1 | Peroxidase | extracellular | redox | 2.69 | 5 | 13 | 21 | NA |
| TraesCS1B01G013500.1 | LMW-GS | vacuole | storage protein | 2.46 | 5 | 7 | 13 | NA |
| TraesCS4A01G112300.1 | GILT | extracellular | not assigned | 2.35 | 5 | 16 | 27 | NA |
| TraesCS1D01G317300.1 | HMW-GS | vacuole | storage protein | 2.21 | 3 | 4 | 5 | NA |
| TraesCS6D01G356600.1 | Aspartic proteinase | golgi | protein degradation | 2.19 | 9 | 37 | 57 | NA |
| **20 most rapidly degraded proteins** |  |  |  |  |  |  |  |  |
| TraesCS6D01G000200.1 | Alpha-AI | golgi | stress response | 0.94 | 3 | 11 | 16 | 0.74 |
| TraesCS4D01G212300.1 | HSP | cytosol | chaperone | 0.92 | 3 | 3 | 4 | 0.75 |
| TraesCS2A01G310300.2 | SBE | NA | major CHO metabolism | 0.91 | 3 | 4 | 4 | 0.76 |
| TraesCS7A01G035600.1 | Gamma-gliadin | vacuole | storage protein | 0.85 | 3 | 3 | 4 | 0.81 |
| TraesCS6D01G089200.1 | Endoglucanase | golgi | misc | 0.82 | 4 | 9 | 10 | 0.84 |
| TraesCS6D01G230000.1 | SAM | cytosol | AA synthesis | 0.82 | 3 | 13 | 16 | 0.85 |
| TraesCS1A01G033600.1 | TIM17 | mitochondrion | protein targeting | 0.8 | 3 | 6 | 11 | 0.86 |
| TraesCS7D01G307500.1 | UGE | golgi | cell wall relative | 0.78 | 3 | 4 | 6 | 0.89 |
| TraesCS4A01G250800.1 | ERO1 | golgi | protein targeting | 0.77 | 3 | 11 | 13 | 0.9 |
| TraesCS7D01G031700.1 | Gliadin-like avenin | vacuole | storage protein | 0.74 | 6 | 14 | 32 | 0.94 |
| TraesCS4D01G252100.1 | Beta-xylosidase | golgi | cell wall relative | 0.73 | 3 | 6 | 7 | 0.94 |
| TraesCS4A01G406200.1 | ADH | cytosol | fermentation | 0.73 | 3 | 9 | 9 | 0.95 |
| TraesCS5D01G135900.1 | UTase | NA | AA metabolism | 0.72 | 3 | 4 | 5 | 0.97 |
| TraesCS4D01G250000.1 | Alpha-AI | golgi | stress response | 0.71 | 3 | 12 | 16 | 0.98 |
| TraesCS1D01G001000.1 | Gamma-gliadin | NA | storage protein | 0.7 | 5 | 5 | 6 | 0.99 |
| TraesCS5D01G004000.1 | GSP | golgi | storage protein | 0.69 | 3 | 4 | 6 | 1 |
| TraesCS7D01G364600.1 | ABA receptor | cytosol | stress response | 0.69 | 3 | 4 | 4 | 1.01 |
| TraesCS5D01G210000.1 | TPS | cytosol | minor CHO metabolism | 0.68 | 3 | 5 | 5 | 1.01 |
| TraesCS1A01G007400.1 | Gamma-gliadin | vacuole | storage protein | 0.68 | 7 | 7 | 19 | 1.02 |
| TraesCS7D01G031800.1 | Gliadin-like avenin | vacuole | storage protein | 0.67 | 4 | 9 | 24 | 1.04 |

The first wheat genome Traes ID number of the protein group, short protein name, subcellular location (from cropPAL.org) and functional category (MapMan) are included. The number of biological replicates (Brep No.), number of unique peptides (Pep No.) and number of mass spectra hits (Spectra No.) for each protein are also shown.

**Extended Data Table 2.** The 20 wheat grain proteins with the highest ATP cost for biogenesis and maintenance during grain development.

| Accession | Protein Name | Location | Functional Category | ATP Cost ($\mu\text{mol grain}^{-1} \text{d}^{-1}$) | ATP Cost (%) | Brep No. | Pep No. | Spectra No. |
| --- | --- | --- | --- | --- | --- | --- | --- | --- |
| **20 highest ATP cost proteins based on protein synthesis** |  |  |  |  |  |  |  |  |
| TraesCS1A01G066100.1 | Globulin-11S | vacuole | storage protein | 0.928 | 1.663 | 6 | 27 | 49 |
| TraesCS1B01G013500.1 | LMW-GS | vacuole | storage protein | 0.752 | 1.348 | 5 | 7 | 13 |
| TraesCS7D01G031700.1 | Gliadin-like avenin | vacuole | storage protein | 0.546 | 0.979 | 6 | 14 | 32 |
| TraesCS1D01G067100.1 | Globulin-11S | vacuole | storage protein | 0.337 | 0.604 | 5 | 13 | 25 |
| TraesCS5A01G554200.1 | Beta-amylase | golgi | major CHO metabolism | 0.271 | 0.485 | 6 | 14 | 21 |
| TraesCS2A01G168200.1 | Susy | cytosol | major CHO metabolism | 0.256 | 0.46 | 9 | 144 | 358 |
| TraesCS1A01G007400.1 | Gamma-gliadin | vacuole | storage protein | 0.227 | 0.406 | 7 | 7 | 19 |
| TraesCSU01G032700.1 | Beta-amylase | golgi | major CHO metabolism | 0.167 | 0.3 | 8 | 44 | 92 |
| TraesCS4A01G296000.1 | Globulin 1 | vacuole | storage protein | 0.167 | 0.299 | 4 | 6 | 8 |
| TraesCS1B01G449700.1 | AGPase | plastid | major CHO metabolism | 0.145 | 0.259 | 7 | 73 | 121 |
| TraesCS4A01G214200.1 | PDI | endoplasmic reticulum | redox | 0.119 | 0.213 | 9 | 214 | 400 |
| TraesCS7B01G072000.1 | Alpah-AI | extracellular | stress response | 0.119 | 0.213 | 7 | 8 | 11 |
| TraesCS5D01G368900.1 | Serpin | golgi | protein degradation | 0.111 | 0.199 | 3 | 14 | 20 |
| TraesCS1B01G330000.1 | HMW-GS | vacuole | storage protein | 0.107 | 0.192 | 7 | 21 | 32 |
| TraesCS4D01G192900.1 | GILT | extracellular | not assigned | 0.102 | 0.183 | 6 | 8 | 10 |
| TraesCS6A01G372500.1 | Aspartic proteinase | golgi | protein degradation | 0.096 | 0.172 | 5 | 9 | 12 |
| TraesCS7D01G284900.2 | AGPase | plastid | major CHO metabolism | 0.094 | 0.168 | 4 | 4 | 5 |
| TraesCS1B01G102700.1 | Aminotransferase | plastid | AA synthesis | 0.094 | 0.168 | 9 | 114 | 175 |
| TraesCS7D01G535400.1 | SBE | plastid | major CHO metabolism | 0.088 | 0.158 | 4 | 11 | 11 |
| TraesCS7D01G031800.1 | Gliadin-like avenin | vacuole | storage protein | 0.079 | 0.142 | 4 | 9 | 24 |
| **20 highest ATP cost proteins based on protein degradation** |  |  |  |  |  |  |  |  |
| TraesCS1B01G330000.1 | HMW-GS | vacuole | storage protein | 0.0585 | 0.105 | 7 | 21 | 32 |
| TraesCS7D01G031800.1 | Gliadin-like avenin | vacuole | storage protein | 0.0433 | 0.078 | 4 | 9 | 24 |
| TraesCS1B01G449700.1 | AGPase | plastid | major CHO metabolism | 0.0244 | 0.044 | 7 | 73 | 121 |
| TraesCS5A01G554200.1 | Beta-amylase | golgi | major CHO metabolism | 0.0242 | 0.043 | 6 | 14 | 21 |
| TraesCS1B01G013500.1 | LMW-GS | vacuole | storage protein | 0.0236 | 0.042 | 5 | 7 | 13 |
| TraesCS1A01G007400.1 | Gamma-gliadin | vacuole | storage protein | 0.0233 | 0.042 | 7 | 7 | 19 |
| TraesCS7D01G168000.1 | Alpah-AI | extracellular | stress response | 0.0209 | 0.038 | 6 | 11 | 22 |
| TraesCS7D01G284900.2 | AGPase | plastid | major CHO metabolism | 0.0205 | 0.037 | 4 | 4 | 5 |
| TraesCS1B01G294300.1 | HSP70 | cytosol | chaperone | 0.017 | 0.031 | 7 | 18 | 32 |
| TraesCS1A01G066100.1 | Globulin-11S | vacuole | storage protein | 0.0167 | 0.03 | 6 | 27 | 49 |
| TraesCS7D01G031700.1 | Gliadin-like avenin | vacuole | storage protein | 0.0145 | 0.026 | 6 | 14 | 32 |
| TraesCS5D01G268000.1 | HSP90 | cytosol | chaperone | 0.0137 | 0.025 | 9 | 234 | 432 |
| TraesCS5D01G020100.1 | Enolase | cytosol | glycolysis | 0.0113 | 0.02 | 9 | 14 | 28 |
| TraesCS2A01G385900.1 | VPE | vacuole | protein degradation | 0.0091 | 0.016 | 3 | 6 | 8 |
| TraesCS3B01G186100.1 | RBCL | plastid | photosynthesis | 0.0091 | 0.016 | 7 | 10 | 17 |
| TraesCS4D01G267600.2 | CDC48-like | cytosol | cell function | 0.0085 | 0.015 | 6 | 6 | 11 |
| TraesCS1B01G443100.1 | GTP-binding protein | cytosol | signalling | 0.0084 | 0.015 | 6 | 12 | 19 |
| TraesCS4D01G250000.1 | Alpha-AI | golgi | stress response | 0.0083 | 0.015 | 3 | 12 | 16 |
| TraesCS4A01G281500.1 | SND1 | cytosol | RNA regulation | 0.0077 | 0.014 | 9 | 66 | 93 |
| TraesCS7D01G454200.1 | Beta-Tubulin | cytosol | cell function | 0.0076 | 0.014 | 9 | 121 | 260 |

The wheat genome first Traes ID of the protein group, short protein name, subcellular location (from cropPAL.org) and functional category (from MapMan) are included. The number of biological replicates (Brep No.), number of unique peptides (Pep No.) and number of mass spectra hits (Spectra No.) for each protein are also shown. ATP Cost (%) column shows the proportion of ATP energy cost to total ATP production ( $55.77 \mu\text{mol grain}^{-1} \text{d}^{-1}$ ) based on respiratory rate measurements.

## Extended Data Figures

**Extended Data Fig. 1.**
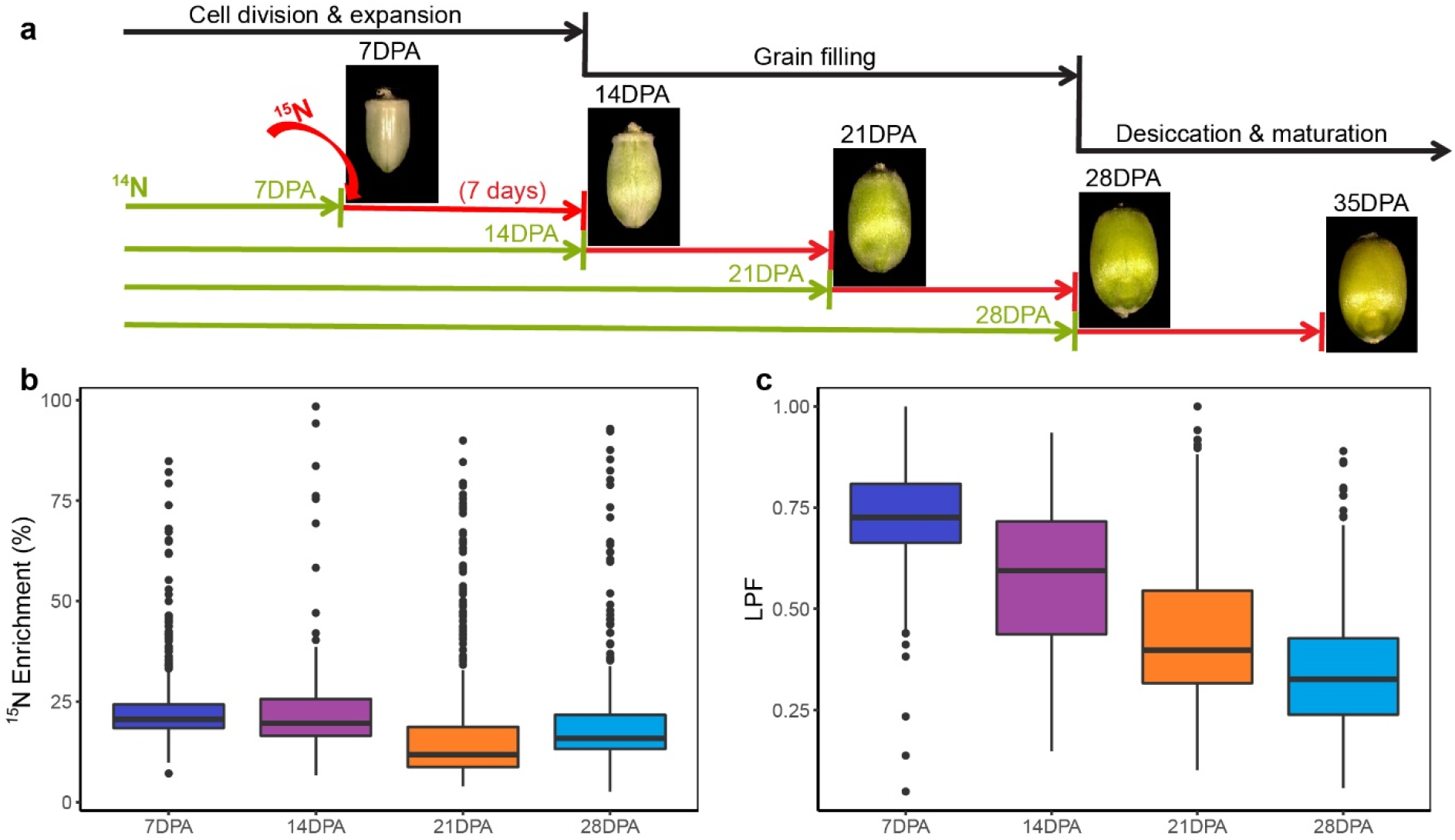
Optimisation of *in vivo* grain protein labelling approach during grain development. **a**, The nitrogen stable-isotope ^15^N was used in this research and provided to plants as N-salts in hydroponic solutions. A fixed period of 7 days of continuous labelling was performed but was started at 7, 14, 21 or 28 days post anthesis (DPA), respectively. A representative grain image from each time point and a brief grain growth stage description is shown. **b**, The degree of nitrogen incorporation in the labelled peptides after 7 days of labelling at each growth stage is shown as ^15^N enrichment level. **c**, The labelled protein fraction (LFP) showing the ratio of the abundance of ^15^N labelled protein to total peptide abundance (natural abundance protein + ^15^N-labelled protein) after 7 days of labelling at each growth stage.

**Extended Data Fig. 2.**
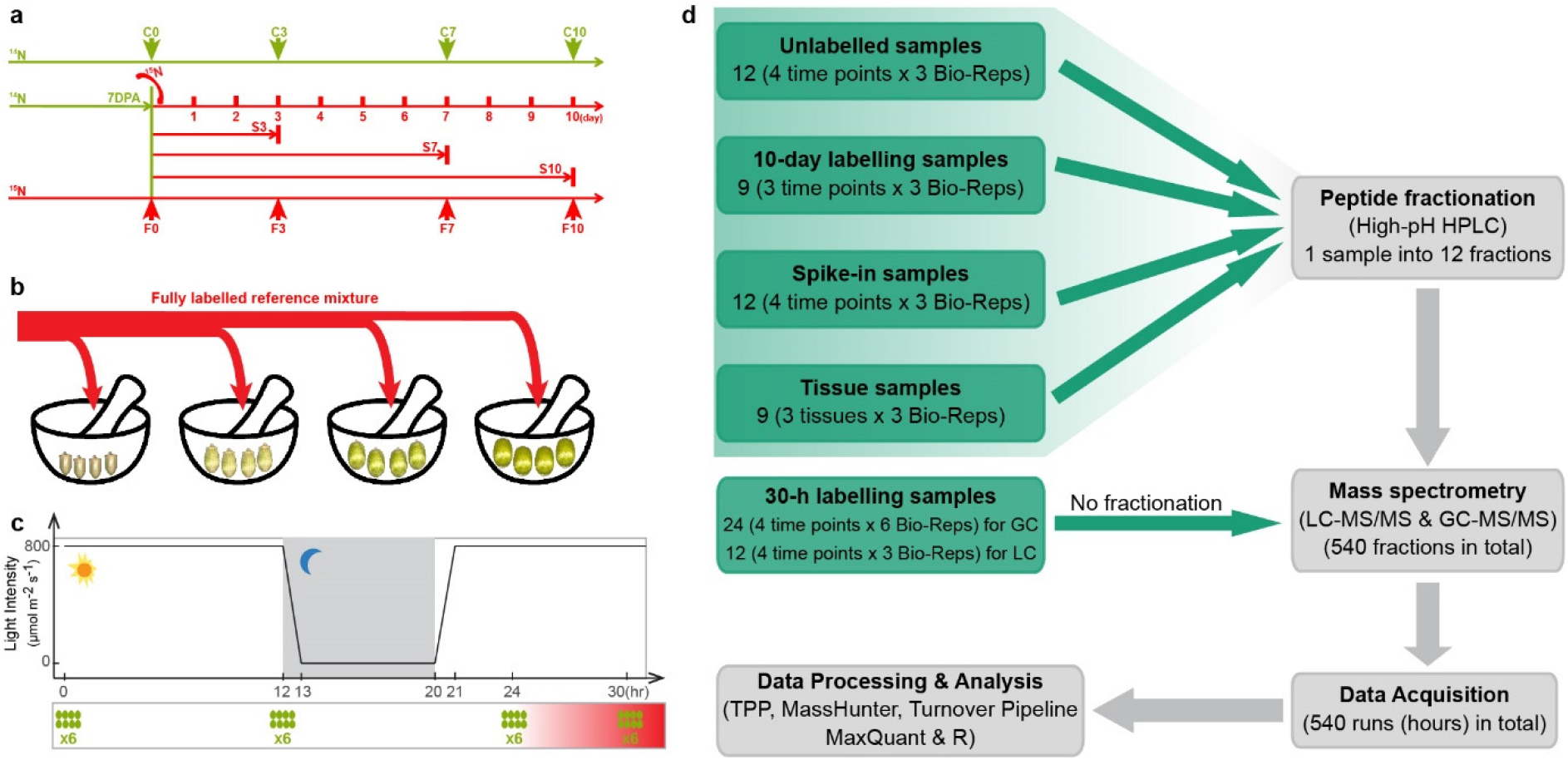
A flow diagram of experiment design, sample preparation, MS data acquisition and analysis. **a**, 10-day progressive labelling experiment for grain protein turnover rates measurement. Labelling started at 7DPA, and samples were collected after 3 days (S3), 7 days (S7) and 10 days (S10) of ^15^N labelling. The unlabelled samples (C0 to C10) and fully labelled samples (F0 to F10, internal reference) at corresponding time points were also collected. Eight grains from the middle of an ear were collected as a biological replicate, and in total three biological replicates were harvested. **b**, A spike-in approach was developed to measure individual wheat grain protein FCP during grain development. The same amount of a fully labelled reference mixture (100 mg in this study) was spiked into four grains at each time point, where the reference mixture powder was prepared by pooling together eight fully labelled grains of each time point. **c**, 30-h progressive labelling experiment for lag time estimation. Using 7DPA plants, samples were collected at 0, 12, 24, 30 hours after labelling, and 8 grains from the middle of an ear were collected as a biological replicate and six biological replicates were collected. **d**, a flow diagram of sample preparation, mass spectrometry data acquisition and afterward data analysis. The 12 30-h progressive labelling samples were used for metabolite measurements by GC-MS/MS and peptide measurements by LC-MS/MS.

**Extended Data Fig. 3.**
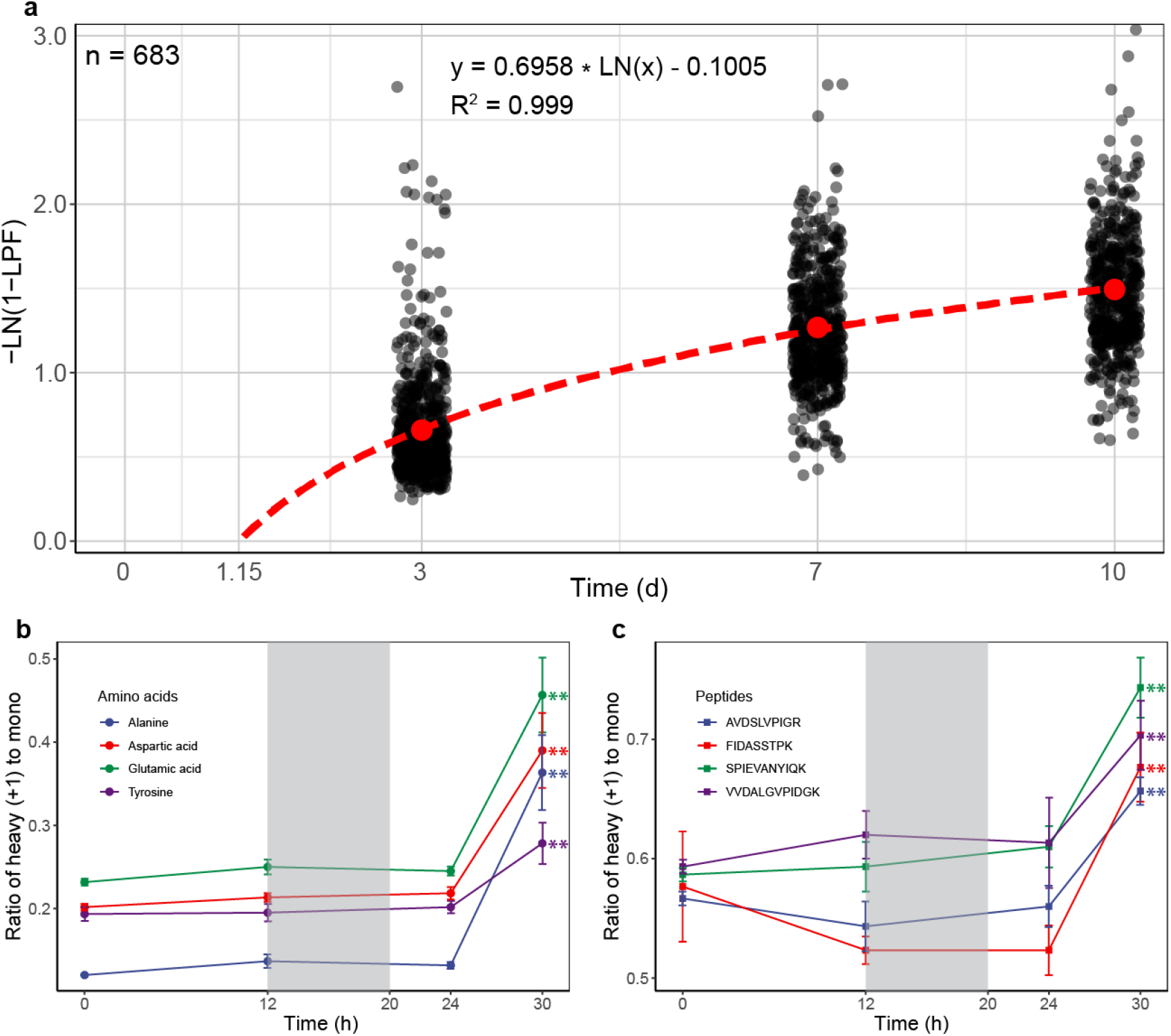
The time lag of label incorporation into wheat grain proteins. **a**, The estimation of lag time of incorporation into proteins via logarithmic regression of the inverse of the natural log of natural abundance protein (1-LPF). The dashed red line demonstrated the logarithmic regression model using the mean value (red dots) of each time point. The formula of the logarithmic regression model and its R-square are shown on the top of the scatter plot. **b**, The ratio of heavy (+1) to mono abundance over the time course of 4 example amino acids measured by GC-MS. Six biological replicates were conducted. **c**, The ratio of heavy (+1) to mono abundance over time course of 4 example peptides measured by LC-MS/MS. Three biological replicates, three of the six used for the GC analysis, were conducted. The shade area represents night time. One-way ANOVA p-value < 0.01 (asterisks). Actual mass spectrum peaks of the representative amino acid and peptide are shown in Extended Data Fig. 4. Detailed data and statistical test results involved in this analysis are listed in Supplementary Data 2.

**Extended Data Fig. 4.**
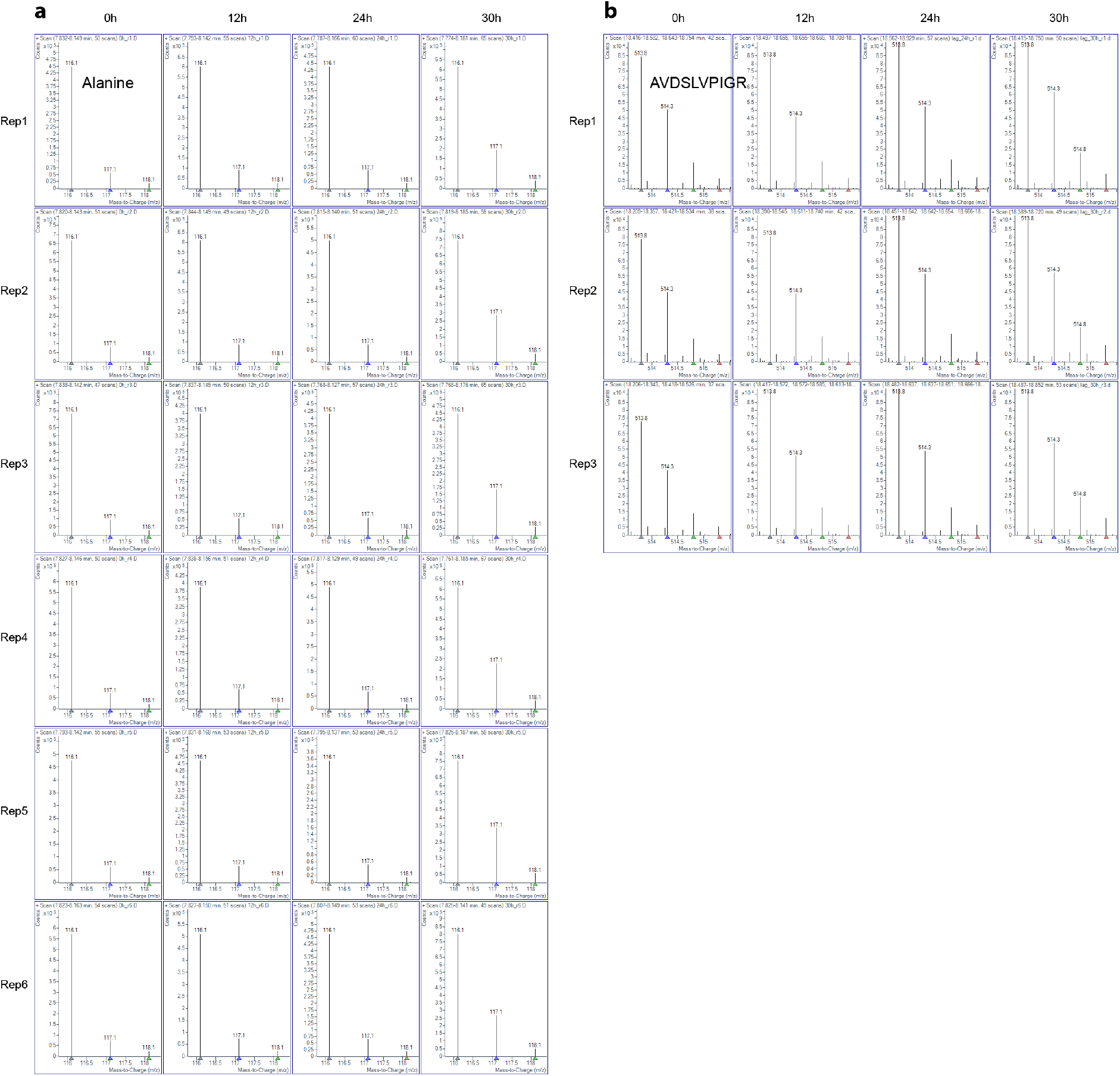
Mass spectrum of the change from mono to mono + labelled abundances of amino acids and peptides over a 30-h time course to estimate labelling lag time. **a**, Example mass spectrum peaks of mono and mono + labelled abundance of alanine over the 30h time course. Unlabelled m/z: 161.1; +1 m/z: 117.1. **b**, Mass spectrum peaks of mono and mono + labelled abundance of the representative peptide (AVDSLVPIGR) over the 30h time course. Unlabelled m/z: 513.8; +1 m/z: 514.3, +2 m/z: 514.8 and +3 m/z: 515.3.

**Extended Data Fig. 5.**
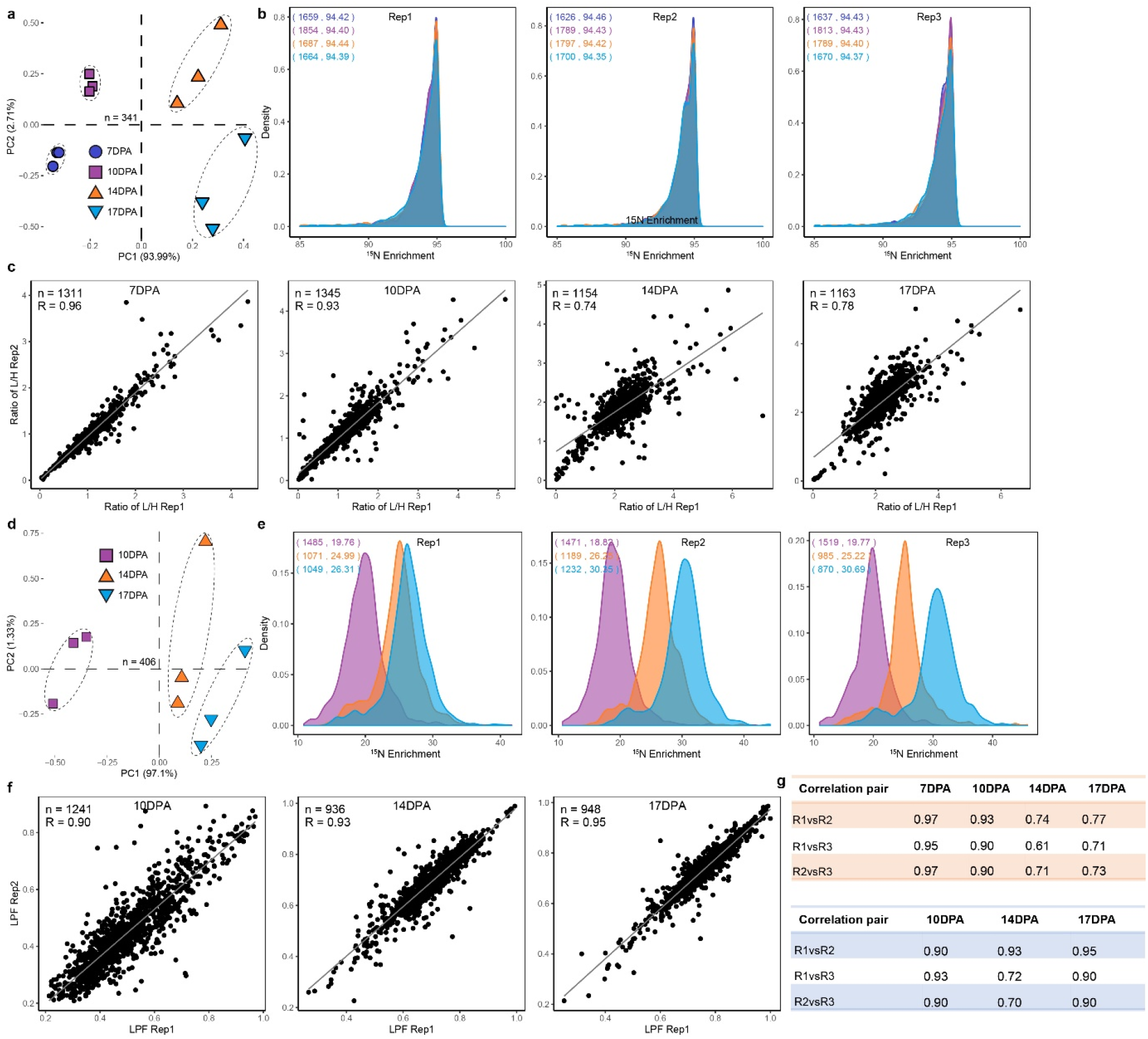
Data quality verification of spike-in and progressive labelling data. A series of data quality verifications were conducted before further downstream data analysis. The raw data used for this analysis are provided in Supplementary Data 3. The high quality of raw data was supported by three factors, namely the high correlation coefficient (0.84 average) between biological replicates; clear separation of different time point samples in PCA; and consistent nitrogen incorporation (^15^N enrichment) between biological replicates at a timepoint. **a**, Principal component analysis of spike-in data. The dashed circles outline biological replicates for each time point. **b**, The ^15^N enrichment level of spike-in data of each time point shown in overlapped density plots. The number of proteins and median of ^15^N enrichment of each time point are shown within brackets (7DPA: indigo; 10DPA: purple; 14DPA: orange; 17DPA: turquoise). **c**, Representative scatter plot showing Pearson correlation coefficient between biological replicate 1 and 2 of spike-in data in light to heavy ratio at each time point. The solid grey line indicates the linear regression model. Data size in number of proteins and the correlation coefficient shown on the top-left corner of the plot. Correlation coefficient of all possible pairs are shown in the upper panel in **g**. **d**, **e** and **f** showing data verification result of progressive labelling data using the same analysis strategies as in A, B and C respectively. G, The Pearson correlation coefficient of all possible pairs of spike-in (upper panel) and progressive labelling data (lower panel).

**Extended Data Fig. 6.**
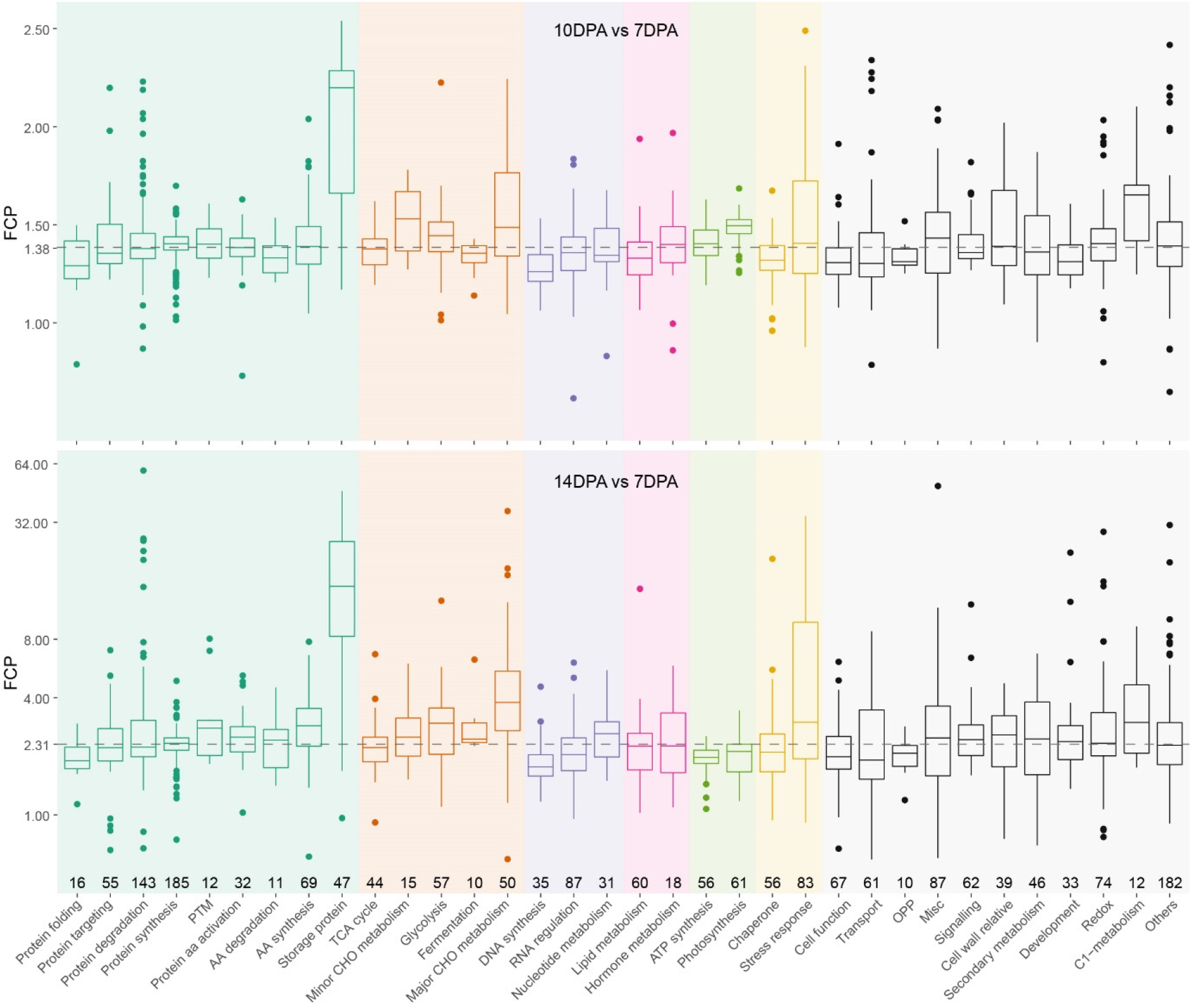
The individual FCP of wheat grain proteins at DPA 10 and DPA 14 summarized by their functional categories. 34 categories (≥ 10 proteins) were displayed, and the rest of the categories (< 10 proteins) were grouped into ‘Others’. The number of proteins for each functional category (MapMan bin) is displayed along with x-axis, and boxes within each superior category are sorted by increasing order of median FCP. The y-axis is log2 transformed, and the dashed line shows the overall median FCP. Pairwise t-test results between categories are listed on Supplementary Data 4b. Colours represent to broad categories: Amino acid metabolism (Green), Carbohydrate metabolism (Orange-red), Nucleotide metabolism (Purple), Lipid metabolism (Pink), Energy producing (Lawngreen), Stress response (Gold) and ‘Other’ categories (Black).

**Extended Data Fig. 7.**
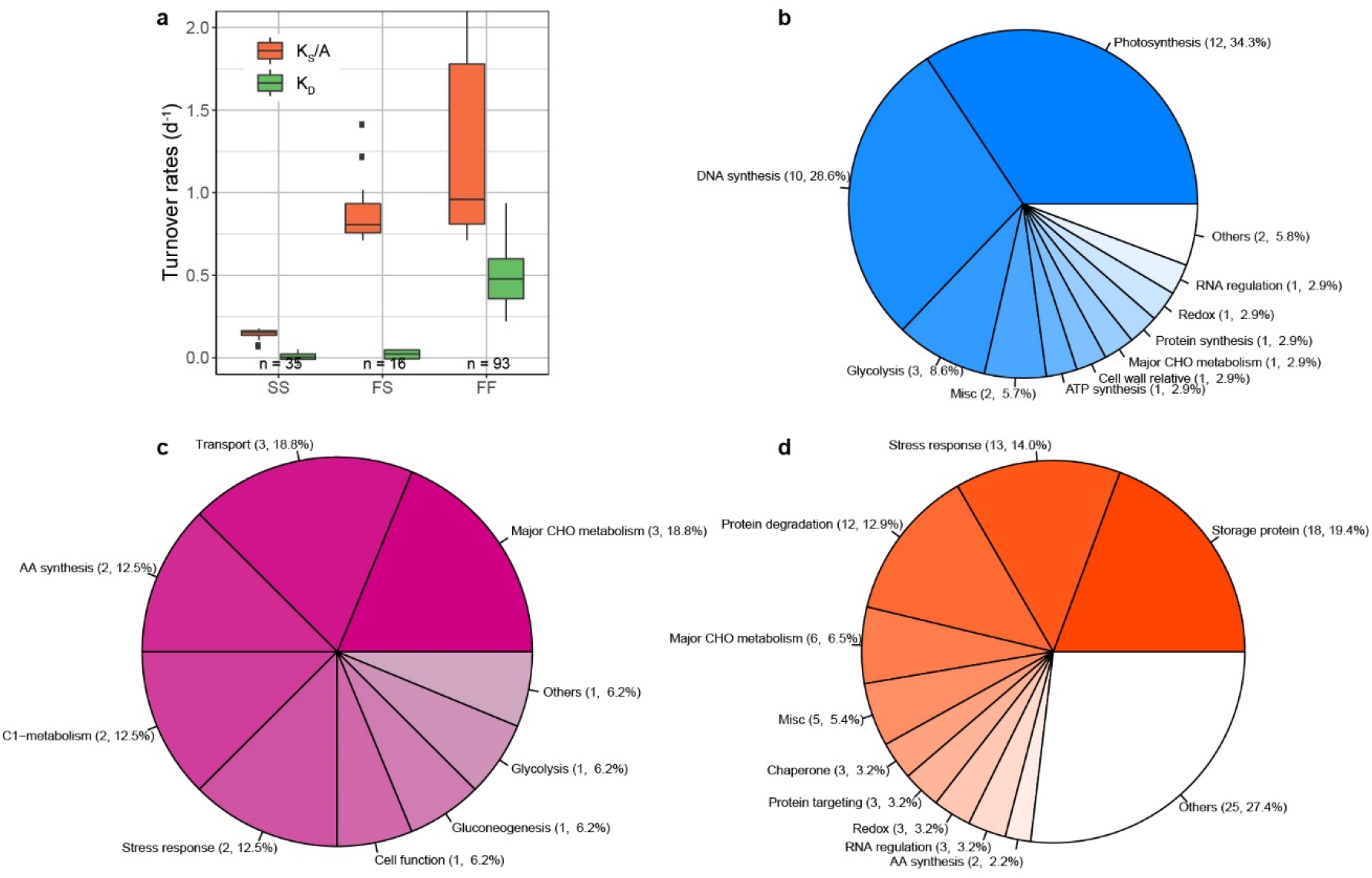
Relatively slow and fast turning over wheat grain proteins during grain development. A total of 678 proteins, defined as relatively fast or slow turning over proteins, were identified within the dataset (Supplementary Data 5b). These proteins were further grouped into three categories according to their turnover characteristics, house-keeping proteins (SS: having both relatively slow K_S_/A and K_D_ rates), induced but stable proteins (FS: having relatively fast K_S_/A but relatively slow K_D_) and rapidly-cycling proteins (FF: having both relative fast K_S_/A and K_D_ rates). **a**, Box plots of the averaged K_S_/A and K_D_ rates calculated over three time points for the SS, FS and FF categories. The number of proteins in each category are shown along with x-axis. Statistical results derived from one-way ANOVA and Tukey’s HSD were shown in Supplementary Data 5c. **b**, Pie plot of functional categories of house-keeping proteins (SS). **c**, Pie plot of functional categories of induced but stable proteins (FS). **d**, Pie plot of functional categories of rapidly-cycling (FF). Only top 10 categories are shown if total functional categories ≥ 10, and the rest of categories are assigned into Others.

**Extended Data Fig. 8.**
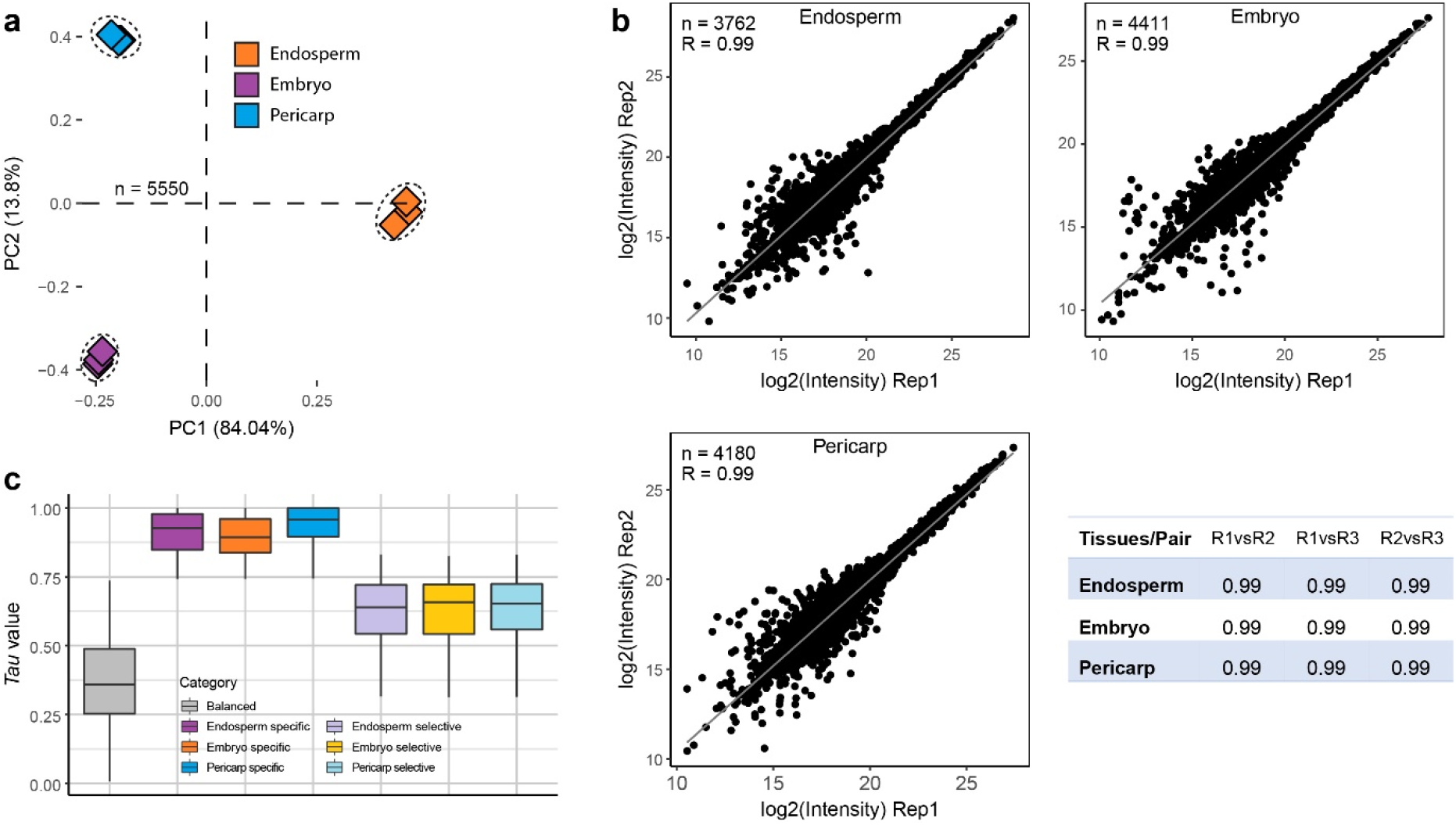
Data quality verification of wheat grain tissue data sets. **a**, Principal component analysis of 5550 protein abundances used for this analysis (Supplementary Data 6). Dashed circles encompass three biological replicates for each time point. **b**, Representative scatter plots showing Pearson correlation coefficient between biological replicate 1 and 2 of log2 transformed protein abundance intensity for each tissue. The solid grey line shows the linear regression model. The number of proteins identified and the correlation coefficient between replicates is shown on the top-left corner of the plot. The table shows correlation coefficients of all possible pairs. **c**, The box plot of *Tau* value for the seven protein expression categories.

**Extended Data Fig. 9.**
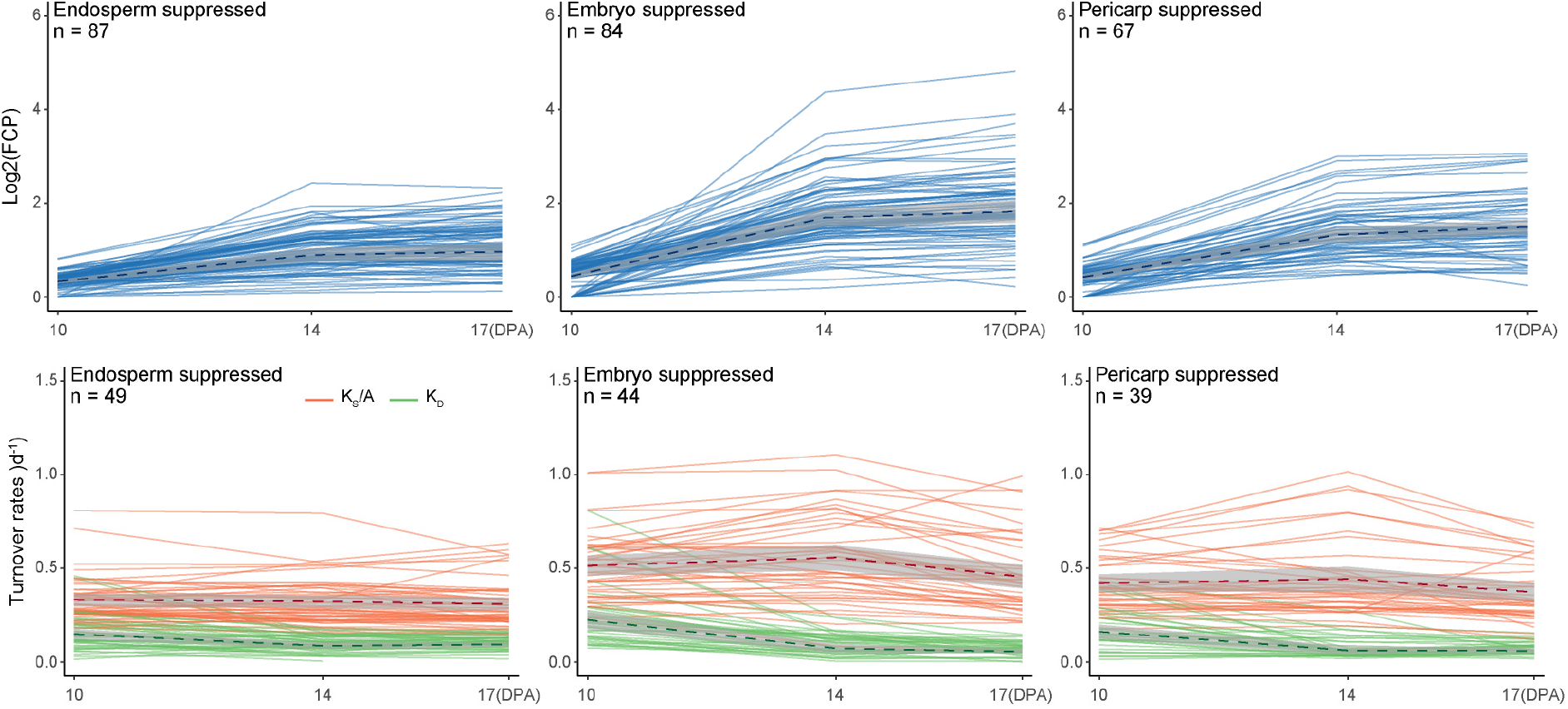
Changes in FCP and turnover rate profiles of proteins present in a lower abundance in a given tissue type during grain development. Only proteins having values (FCP or turnover rates) at all three time points are included in the analysis. The dashed lines demonstrate the mean values, and the grey shade areas show the 95% confidence intervals.

**Extended Data Fig. 10.**
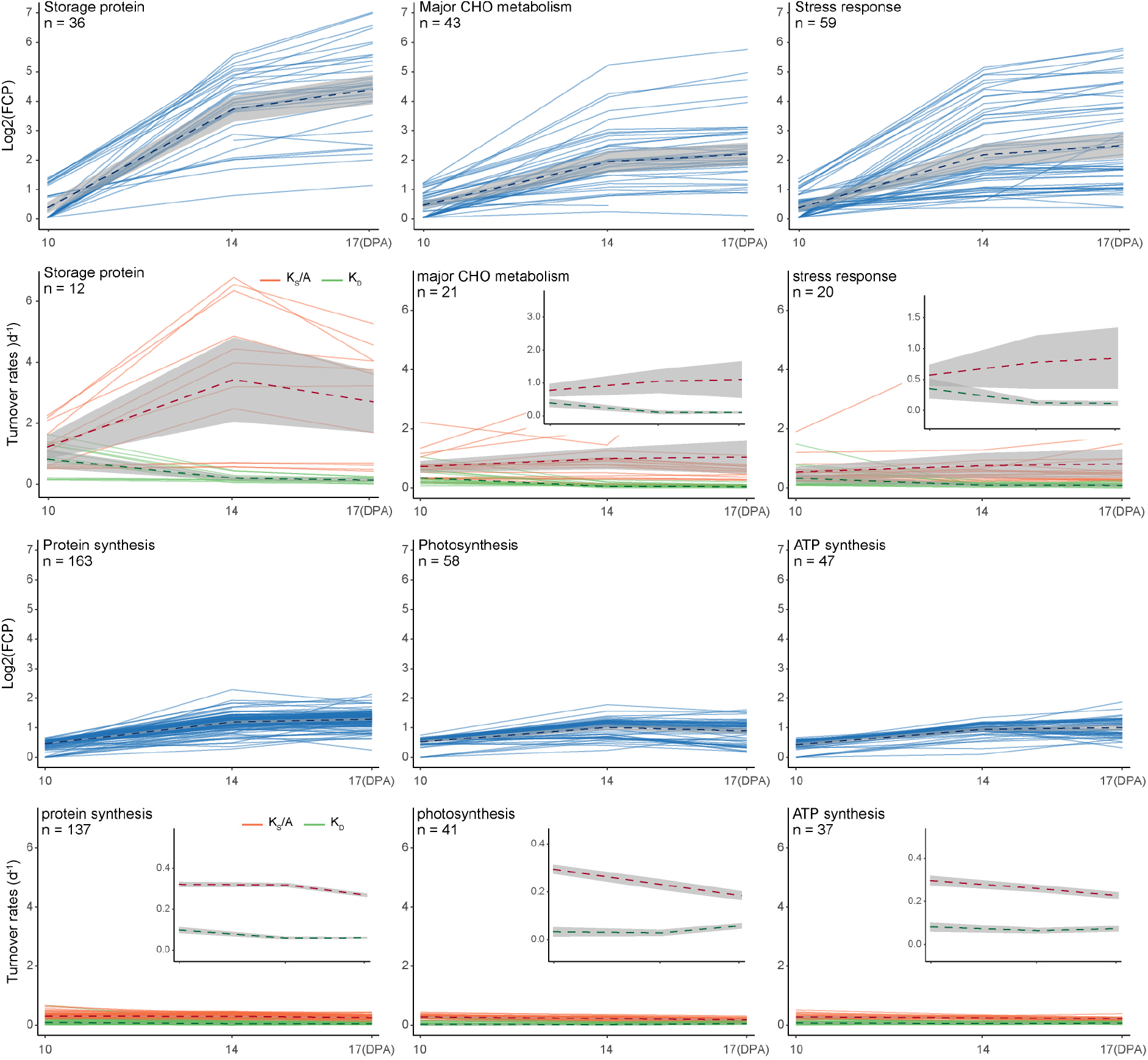
Changes in turnover rates of proteins with high (top 6 graphs) and low (bottom 6 graphs) FCP during grain development. Each pattern includes three example functional categories. FCP (Blue), protein synthesis K_A_/S (Orange) and protein degradation K_D_ (Green) values are shown. In each functional category, only proteins with FCP or turnover rates measured at all three time points are included. The number of proteins included in each functional category is shown. Dashed lines indicate the mean values, and the grey shade areas show the 95% confidence intervals. The optimized y-axis scale version of turnover rates data are inserted to highlight the change pattern.

## Supplementary information

**Supplementary Data 1**

Full data list of grain fresh weight, grain respiration rate and total grain ATP production.

**Supplementary Data 2**

Full data list of lag time modelling, GC-MS/MS data and LC-MS/MS data.

**Supplementary Data 3**

Full data list used for data quality control analysis.

**Supplementary Data 4**

Full data list of wheat grain individual FCP values during grain development.

**Supplementary Data 5**

Full data list of wheat grain protein turnover rates during grain development.

**Supplementary Data 6**

Full data list of protein relative abundance for embryo, endosperm and pericarp proteomes.

**Supplementary Data 7**

Full data list of ATP energy budget used for wheat grain protein turnover during grain development.

**Supplementary Data 8**

Full data list of wheat grain key storage proteins accumulation profiles during grain development.

